# Structural basis of the human transcriptional Mediator complex modulated by its dissociable Kinase module

**DOI:** 10.1101/2024.07.01.601608

**Authors:** Shin-Fu Chen, Ti-Chun Chao, Hee Jong Kim, Hui-Chi Tang, Subash Khadka, Tao Li, Dung-Fang Lee, Kenji Murakami, Thomas G. Boyer, Kuang-Lei Tsai

## Abstract

The eukaryotic Mediator, comprising a large Core (cMED) and a dissociable CDK8 kinase module (CKM), regulates RNA Polymerase II (Pol II)-dependent transcription. cMED recruits Pol II and promotes pre-initiation complex (PIC) formation in a manner inhibited by the CKM, which is also implicated in post-initiation control of gene expression. Herein we report cryo-electron microscopy structures of the human complete Mediator and its CKM, which explains the basis for CKM inhibition of cMED-activated transcription. The CKM binds to cMED through an intrinsically disordered region (IDR) in MED13 and HEAT repeats in MED12. The CKM inhibits transcription by allocating its MED13 IDR to occlude binding of Pol II and MED26 to cMED and further obstructing cMED-PIC assembly through steric hindrance with TFIIH and the +1 nucleosome. Notably, MED12 binds to the cMED Hook, positioning CDK8 downstream of the transcription start site, which sheds new light on its stimulatory function in post-initiation events.

## Introduction

The eukaryotic Mediator complex is a multisubunit (25 to 30 proteins), evolutionarily conserved transcriptional coactivator that transmits regulatory signals from activators and repressors to the RNA polymerase II (Pol II) transcription machinery^1–3^. Given its large size, inherent kinase activity, and multiple subunit-specific protein interactions, Mediator is not only involved in the assembly of the preinitiation complex (PIC) at promoters during transcription initiation but also participates in regulating Pol II pause and release prior to productive elongation^4,5^.

Structurally, the Mediator complex is composed of a large Core (cMED), consisting of Head, Middle, and Tail modules, as well as a dissociable CDK-containing subcomplex, known as the CDK8 kinase module (CKM) (Figure 1A)^6–8^. Functionally, cMED stimulates basal transcription by recruiting Pol II to promoters, stabilizing PIC assembly, and stimulating the phosphorylation of the Pol II C-terminal domain (CTD) by TFIIH ^3,9^. In metazoans, cMED includes an additional subunit, MED26, that participates in recruiting either TFIID or Super elongation complex (SEC) to promoters for gene activation^10,11^. The CKM is a four-subunit complex, comprising MED12, MED13, CCNC, and CDK8, which can reversibly associate with cMED through its MED13 subunit^7,8,12^. In humans, MED12 has been demonstrated to directly interact with and functionally stimulate the enzymatic activity of CDK8, a colorectal cancer oncogene^13–15^. In vertebrates, three sequence-distinct paralogous subunit pairs within the CKM have emerged, including MED12/12L, MED13/13L, and CDK8/CDK19; together with CCNC, the paralogous subunits assemble into the CKM in a mutually exclusive manner^16^. Based on its abilities to inhibit cMED-activated transcription *in vitro* and preclude interaction of Pol II with cMED, the CKM was initially considered a repressor of transcription^6,7,17^. Recent studies, however, have provided definitive evidence that the CKM also plays a positive role in gene expression by participating in the regulation of Pol II pausing and release during post-initiation events ^18–20^. In this regard, the cMED-CKM has been shown to interact with p-TEFb, which controls the release of paused Pol II ^20^. Additionally, CDK8 phosphorylates several pausing-related factors, and inhibition of its kinase activity results in increased Pol II pausing^21–23^.

**Figure 1.**
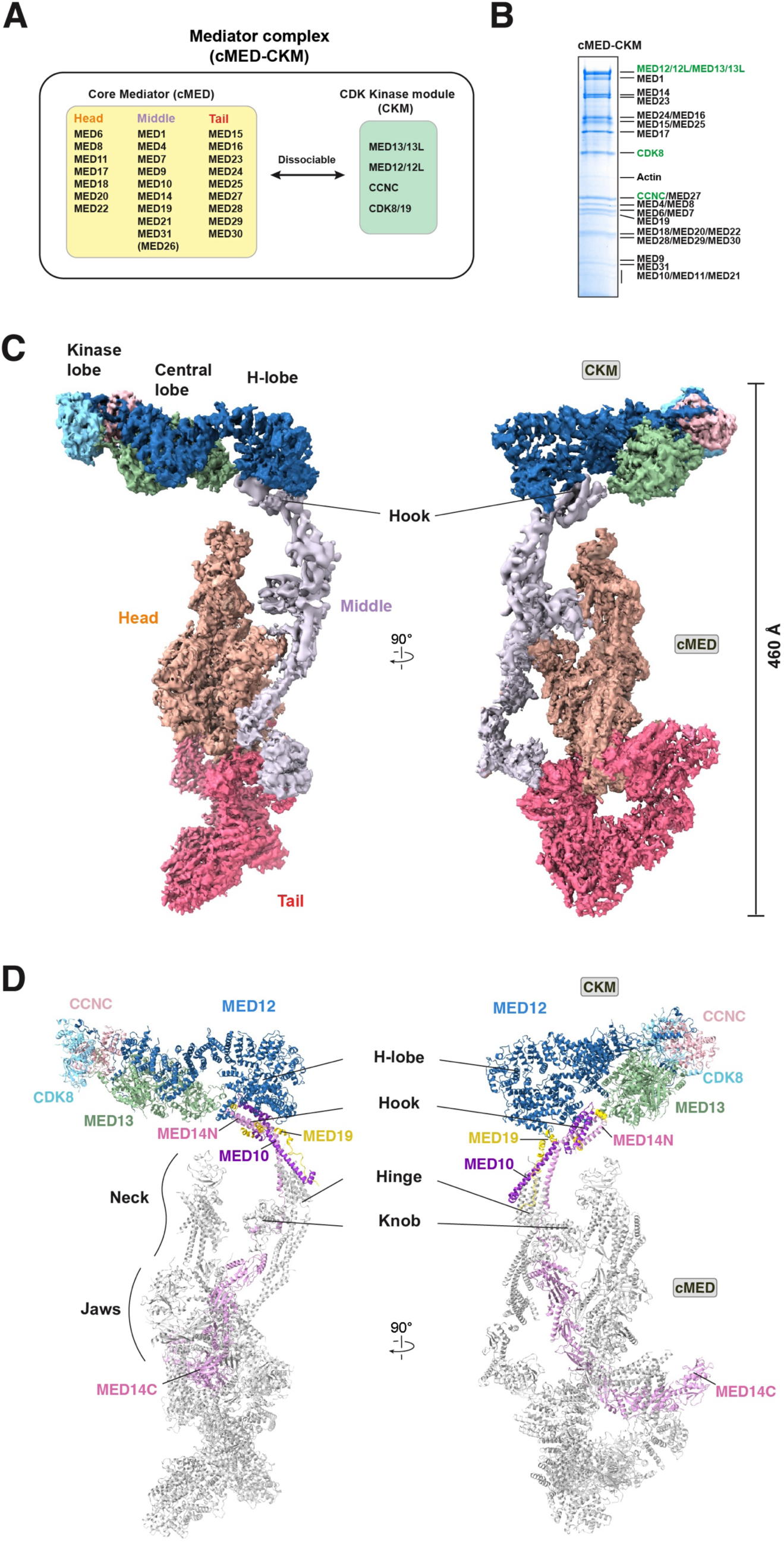
Structure of the human Mediator complex. (A) Subunit composition of the cMED and CKM. In vertebrates, CKM has three paralogous subunit pairs, MED12/12L, MED13/13L, and CDK8/19. The Metazoan-specific subunit, MED26, is indicated by brackets. **(B)** SDS-PAGE of the purified Mediator complex. CKM subunits, including MED12 and MED13 paralogs, are indicated by green text. **(C)** Composite cryo-EM map of human cMED-CKM in two different views. The average resolutions for cMED and CKM are 4.7 Å and 6.7 Å, respectively. The Head, Middle, Tail, and CKM modules as well as other structured regions are labeled as indicated. **(D)** Atomic model of human cMED-CKM in two views, with colored subunits.

Prior structural studies of cMED and its complex with the PIC or PIC-nucleosome have provided mechanistic details into how cMED-PIC assembles with promoter DNA and stimulates CTD phosphorylation via TFIIH to activate transcription ^24–30^. More recently, the structure of yeast CKM established its subunit organization and interactions, the mechanism of Med12-dependent Cdk8 activation, and the architecture of an Argonaute (Ago)-like Med13 that includes a large intrinsically disordered region (IDR)^8,31^. Despite these advancements, high-resolution structural details of the complete human Mediator, comprising cMED bound to the CKM, have not heretofore been reported. Although low-resolution EM reconstructions of human and yeast cMED- CKM complexes were reported^6–8^, little is known about the mechanism by which the CKM represses cMED-dependent transcription, how it enhances gene expression through CDK8 activity, and the functional significance of the MED13 IDR. Importantly, how the CKM inhibits binding of Pol II and metazoan MED26 to cMED remains poorly understood.

Here we present cryo-EM structures of the complete human Mediator complex and its CKM at near-atomic resolution. Combined with biochemical and crosslinking mass spectrometry (XL-MS) analyses, this study provides a structural basis for the Mediator complex in a repressed state, elucidating the mechanism by which the CKM suppresses cMED-driven transcription. We find that the CKM binds to cMED through two distinct interfaces corresponding to the MED13 IDR and MED12 HEAT repeats. By allocating its IDR to their respective binding sites, the CKM inhibits the binding of Pol II and MED26 to cMED. In addition, the CKM affects cMED-PIC assembly by creating steric hindrance with TFIIH and the proximal +1 nucleosome. Notably, the CKM binds the Hook of cMED in a specific orientation that places CDK8 proximal and downstream relative to the transcription start site (TSS). Finally, we find that human MED12 adopts a conformation distinct from that of yeast MED12 in order to activate CDK8. Altogether, this study reveals how the human CKM represses cMED-activated transcription prior to initiation and highlights its potential contribution to the post-initiation process.

## RESULTS

### Structure determination of human Mediator and its dissociable CKM

We purified endogenous Mediator from 293-F cells expressing Flag-tagged CDK8 through sequential FLAG IP and glycerol gradient centrifugation (Figures S1A and S1B). Peak fractions containing cMED and CKM proteins were pooled and analyzed by SDS-PAGE, mass spectrometry, and *in vitro* kinase assay (Figures 1B and S1C). As expected, the purified complex contained the complete ensemble of Mediator subunits with the exception of MED26 and CDK19 (Data S1). Notably, nearly all particles correspond to the Mediator complex as revealed by negative-stain EM, with little detectable free CKM, suggesting stable interaction and a nearly stoichiometric ratio of cMED to CKM (Figure S1D). Purified Mediator complex was concentrated to ∼ 1 mg/ml and subjected to single-particle cryo-EM data collection using a Titan Krios with K2 direct detector (Figure S1E). 2D class averages calculated from cryo-EM images showed that the CKM binds in one particular orientation relative to cMED (Figures S1F and S1G). Following 2D and 3D classification, ∼105K imaged particles of Mediator were subjected to 3D reconstruction and refinement, resulting in a ∼4.8 Å density map in which apparent cMED secondary structural features were revealed, while the CKM density map was blurred out due to its slight mobility (Figures S2A and S2B). 3D classification was used to obtain an overall density map of the Mediator complex at ∼11 Å, which clearly showed CKM bound to the Hook of cMED (Figure S2C). Next, we divided the Mediator complex into two parts, CKM with the Hook (CKM-Hook), and cMED without the Hook (cMED-HookΔ), for separate data processing. Further focused classification and refinement strategies largely improved resolution and map quality for cMED- HookΔ (4.7 Å) and CKM-Hook (6.7 Å) (Figures S2D and S2E). Both individually refined maps were fitted into the entire Mediator density map and combined to produce a composite map (Figure 1C).

To gain a deeper structural understanding of the human CKM, particularly due to its limited resolution in cMED-CKM, recombinant baculovirus-expressed CKM subunits, including MED13, MED12, CCNC, and CDK8, were co-purified from insect cells via Flag-tagged CDK8, followed by glycerol gradient centrifugation (Figures S1H and S1I)^14^. Purified CKM subunit composition and enzymatic activity were evaluated by SDS-PAGE and kinase assay, respectively (Figures S1C and S1J). Purified enzymatically active CKM was then concentrated to ∼2 mg/ml and subjected to structure determination by single-particle cryo-EM (Figure S1M and table S1). In brief, ∼107,000 particle images were obtained using 2D and 3D classifications, followed by particle polishing and 3D refinement, resulting in an overall 3.8 Å cryo-EM map, in which we noticed that one of the ends (H-lobe, corresponding to the MED12 HEAT repeats) showed poor density due to its mobility (Figures S2H and S2I). Accordingly, to improve map quality, the H-lobe and other regions were further classified and refined to 6.5 and 3.6 angstroms, respectively (Figures S2J and S2K). An atomic CKM model was built into the cryo-EM density maps using the crystal structure of human CDK8/CCNC^32^ and AlphaFold models of MED12 and MED13 as templates (Figure S3). The atomic model of the Mediator complex was built based on global- and focused-refined maps with the structures of human cMED and CKM as templates (Figures 1D and S4 and Table S1).

### Architecture of human Mediator

The structure of the Mediator complex reveals a modular architecture with approximate dimensions of ∼460 Å x 210 Å x 160 Å, which contains all Mediator subunits (excluding MED26 and paralogs of CDK8, MED12, and MED13) that are clearly visualized and organized into a large Core containing Head, Middle, and Tail modules, and an elongated CKM, consisting of Kinase-, Central-, and H-lobes, at the top (Figures 1C and 1D and movie S1). The N-terminal region of MED14 (MED14N) within cMED-CKM adopts an extended conformation comprising several α- helices that run along with the Knob of the cMED Middle module toward the top of cMED, where it interacts with MED10 and MED19 to form the cMED Hook domain (Figure 1D). As revealed by 2D class averages and the 3D structure, the Hook is bound by the CKM with an orientation that is roughly perpendicular to the Middle module (Figures 1D and S1G right). In this regard, the Central- and H-lobes of the CKM form direct contacts with the Hook, while the Kinase-lobe at the opposite end of the CKM is positioned distally and away from the cMED, where it makes no contacts with other modules. Notably, this binding mode is distinct from that described in prior low-resolution EM studies of the yeast Mediator complex, in which the CKM, via multiple contacts, binds to the cMED Middle module in a parallel orientation that overlaps the binding site for Pol II ^8,17^. This suggests that the mechanism by which the CKM inhibits the binding of Pol II to cMED differs between human and yeast (see Discussion). Although the main body of the human CKM primarily utilizes its H-lobe to contact the cMED Hook domain, its orientation relative to cMED nonetheless seems to be quite stable, exhibiting only limited mobility (Figure S2C). This mobility likely derives from flexibility in the Hook domain itself as observed in published structures of cMED^25,26^.

### Architecture of human CKM

Although the CKM in humans shares the same subunit organization as its yeast counterpart, it nonetheless exhibits distinct structural characteristics in MED12 and MED13 that distinguish it from the yeast CKM. The human CKM possesses an elongated architecture with a size of <u>210 × 50 × 85</u> Å (Figure 2A and movie S2). The Kinase-lobe with an ATP-binding site facing outward is mainly comprised of CDK8, CCNC, and two extended regions of MED13 as well as the N-terminus of MED12, while the Central-lobe consists of the MED13 Ago domains and partial MED12 that together encompass a ∼15 Å wide groove. The H-lobe, belonging to MED12, consists of numerous closely packed α-helices and exhibits conformational heterogeneity due to limited mobility. MED13 is the largest subunit of human Mediator (∼250 kDa) and possesses an Ago-like structure comprising four globular domains (N, PAZ, MID, and PIWI) and two linker regions (L1 and L2) that together form two lobes and a self-blocked central channel (Figure 2B). Notably, MED13 harbors a large and unique insertion between the PAZ and L2 domains corresponding to its large IDR (residues 350 – 1069), as well as three additional small insertions, PAZ-ins, PIWI- ins, and MID-ins, that are not present in classical Ago proteins, but all of which are involved in contacts with other CKM subunits (Figure 2B). Unlike the structures of classical Ago proteins in which DNA or RNA are bound in their PAZ and MID domains or central channels, no nucleic acid density was observed in MED13 (Figure S5A). Interestingly, the L2 of human MED13 forms a well-organized structure containing several α-helices that block both nucleic acid binding sites in the PAZ domain and the central channel. Together, these structural observations indicate that MED13 may adopt a different nucleic acid-binding mode than canonical Ago protein family members. Finally, we identified two zinc-binding sites (I and II) present in human, but not yeast, MED13. One of these (site I) is located on the surface between the L2 and PIWI domains and is coordinated by four Cys residues (C1096, C1098, C1122, and C1124) from two loops of L2 (Figure 2C left). The other (site II) is bound by two His residues (H1727 and H1729) of the MED12 C-terminal segment and two Cys residues (C1896 and C1899) of the MED13 PIWI domain near the interface between the Kinase- and Central-lobes (Figure 2C right). These two zinc-binding sites potentially contribute to complex stability or protein-protein interactions, and it is likely that site II is compromised by MED12 mutation H1729N that is causative for Ohdo syndrome, a rare blepharophimosis-associated intellectual disability disorder ^33^.

**Figure 2.**
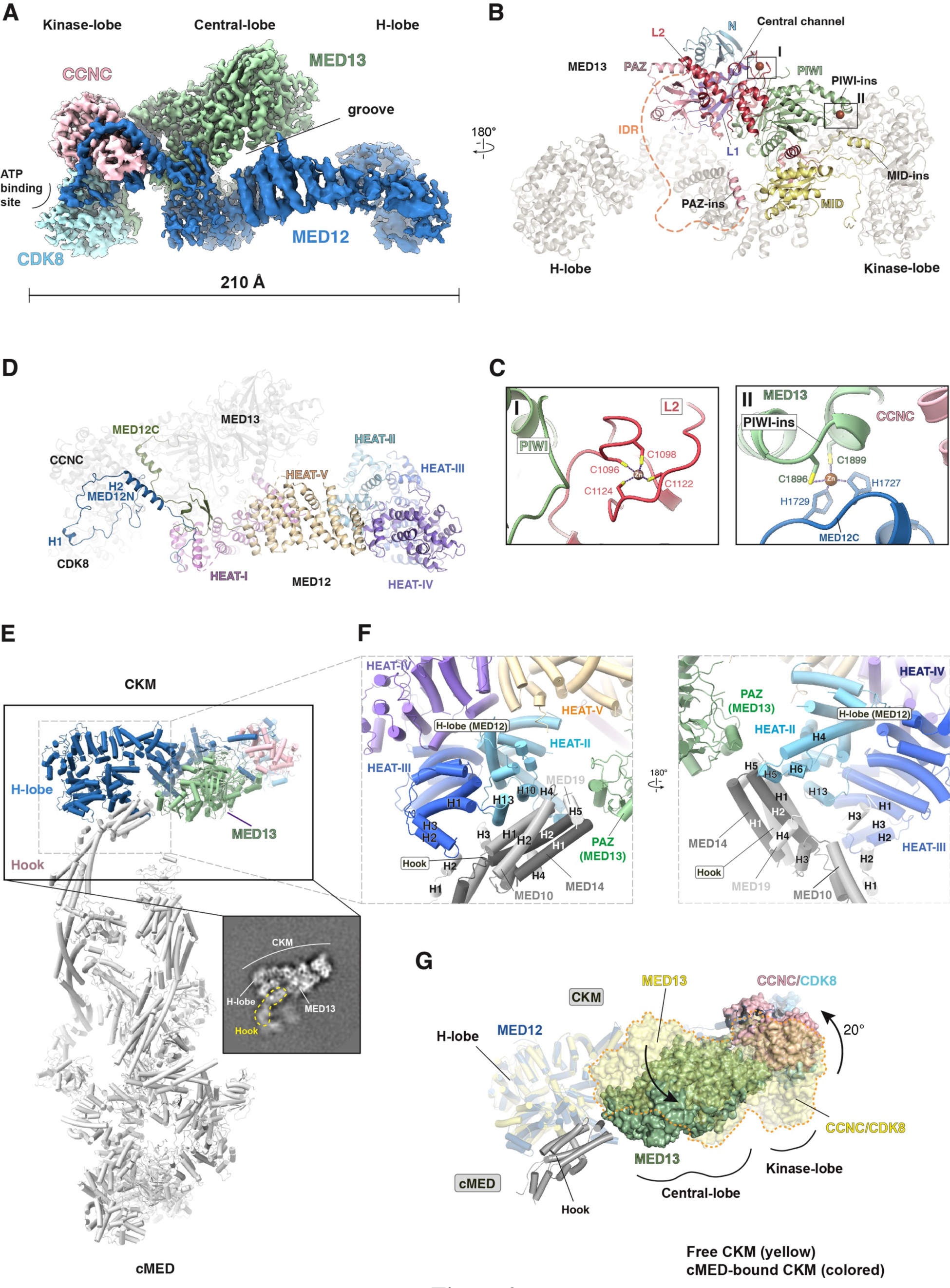
Structure of the human CKM and its interaction with the cMED Hook. (A) Composite cryo-EM map of human CKM. The resolutions of the Kinase-, Central-, and H-lobes are 3.8 Å, 3.6 Å, and 6.5 Å, respectively. **(B)** Structure of MED13 within the human CKM. The positions of Ago (N, PAZ, MID, PIWI) and Linker (L1, L2) domains as well as three insertions (PAZ-ins, MID-ins, PIWI-ins) are indicated. The missing MED13 IDR (residues 350-1069) is indicated by dashed lines. The localization of the Central channel is indicated. Putative Zn metals are indicated by red spheres. Other CKM subunits are colored in gray. **(C)** Close-up views of two zinc-binding sites. Left, Zn^2+^ ion coordinated by 4 Cys residues in the L2 domain. Right, Zn^2+^ ion coordinated by 2 Cys and 2 His residues from MED13 PIWI and MED12C, respectively. The location of both zinc-binding sites (I and II) in MED13 is indicated in (B). **(D)** Structure of MED12 within the CKM. MED12N, MED12C, and five HEAT domains are indicated. Other CKM subunits are colored in gray. **(E)** A 2D class average showing the CKM bound to the cMED Hook. The signals of the CKM-Hook were subtracted from cMED-CKM particles and subjected to 2D classification. The density corresponding to the Hook is encircled by a yellow dashed line. **(F)** Two close-up views of the interface between the cMED Hook and the CKM H-lobe. The α-helices involved in the interaction between cMED and CKM are indicated. **(G)** Superimposition of cMED-bound and free CKM structures by aligning their H-lobes reveals conformational changes in the Kinase- and Central-lobes. The position of MED13 and CDK8/CCNC in free CKM is rendered in transparent yellow.

Both the N- and C-terminal portions of MED12 (residues 1-97 and 1640-1742, hereafter MED12N and MED12C, respectively) adopt extended conformations that are connected by five HEAT-repeat domains (HEAT-I to V) (Figure 2D). MED12N, part of the Kinase-lobe, contains two α-helices that make extensive interactions with CDK8/CCNC. The HEAT-I domain of MED12 lies within the Central-lobe and directly contacts the MID domain of MED13, while the remaining four HEAT domains are tightly packed to form the H-lobe. MED12C runs along the interface between MED13 and HEAT-I, forms an α-helix engaged between the Kinase and Central-lobes, and ultimately binds the MED13 PIWI domain through multiple contacts (Figure 2D). Following MED12C, the PQL domain (residues 1763-2051) that is enriched for proline, glutamine, and leucine amino acids and previously reported to interact with several transcription factors, including Δ-catenin, SOX9, and REST, is conspicuously missing in our density maps^34,35^. Based on our structure, the MED12 N-terminus is consigned to functional interaction with CDK8/CCNC, while the majority of MED12 functions as a structural scaffold with multiple HEAT domains and unstructured regions that offer a large surface area for protein-protein interactions.

### Structural analysis of the cMED-CKM interface

The map density at the interface between cMED and the CKM was initially poor due to flexibility of the cMED Hook. By using focused image analysis, we were able to improve the map quality and visualize the interactions at the interface more clearly (Figures 2E and S2E). Their contacts are confined to a small region, where the Hook is primarily bound by the H-lobe of the CKM and to a lesser extent through its Central-lobe, resulting in a total interaction area of ∼680 Å^2^ (Figures 2E and S4B). At the top of the Hook, H10 and H13 within HEAT-II of the H-lobe contact H1 and H2 of MED10, while H2 of H-lobe HEAT-III interacts with H2 of MED19 (Figure 2F left). Near the tip of the Hook, MED14 H2 is bound by H5 of H-lobe HEAT-II, while MED19 H5 is slightly contacted by the MED13 PAZ domain (Figure 2F). Interestingly, within structure of the free CKM most of the loop densities from the H-lobe involved in contacts with the cMED Hook are missing, suggesting that Hook binding can stabilize these flexible CKM loops. By employing this binding mode, CDK8/CCNC is stably positioned at a distance of ∼120 Å from the Hook without establishing any contacts with cMED. Together, our structure not only explains early biochemical observations that MED12 and MED13 can associate with MED14 and MED19^8^, respectively, but also uncovers a new functional role for the cMED Hook in CKM binding and directional orientation of CDK8. Additionally, our Mediator structure reveals that the CKM binds to the cMED Hook primarily through its MED12 H-lobe, as opposed to MED13 that was previously implicated based on lower resolution EM studies^7,8^.

To determine whether cMED or the CKM undergo structural rearrangements upon their mutual binding, we aligned their unbound and bound structures for comparison. For the CKM, we found that the Kinase- and Central-lobes undergo a ∼20 degree rotation relative to the H-lobe upon CKM interaction with cMED that allows the Hook to be well engaged between the H-lobe and MED13 (Figure 2G). On the other hand, we did not observe any significant structural rearrangements within cMED upon CKM binding. Only subtle conformational changes were noted in the Middle module, which undergoes a counterclockwise rotation of approximately 5 degrees along the Knob (data not shown). Based on these observations, it is likely that the CKM inhibits binding of Pol II to cMED not by causing conformational changes within cMED, but through another mechanism, such as steric hindrance.

### XL-MS confirms the distinct cMED-CKM binding mode and reveals a functional role for the MED13 IDR

To strengthen our structural observations, we performed XL-MS analysis on both purified free CKM and cMED-CKM (Data S2 and S3). For the CKM, 130 unique residue pairs (URPs) were identified, 69 of which were mapped onto the CKM structure with a relatively low violation rate (Cα–Cα distances longer than 35 Å) of 13.1% (Figures 3A, S6A and S6C). These mapped URPs agreed well with the overall architecture and subunit organization of the CKM. The remaining 61 unmapped URPs were localized to unstructured loops or large IDRs. Consistent with the CKM structure, the IDRs of MED12 and MED13, which were missing in the density maps, were largely devoid of intersubunit or intrasubunit crosslinks, except for their N-terminal regions (Figure 3A). These findings suggest that, in the absence of cMED, both IDRs adopt flexible conformations that are exposed to solvent without forming stable interactions with the main CKM body.

**Figure 3.**
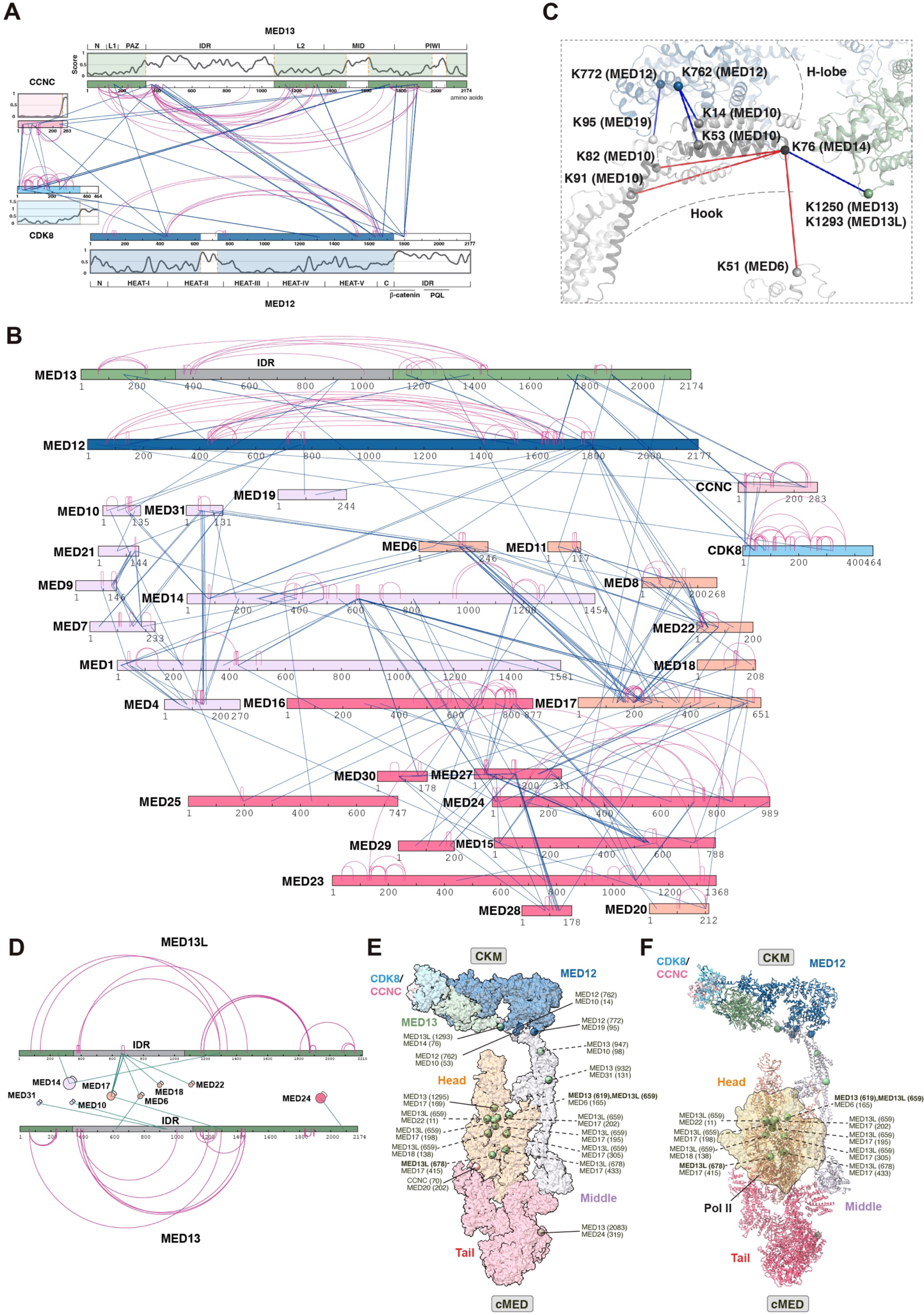
XL-MS analyses of the Mediator complex and its CKM. (A) Crosslinking map and predicted intrinsically unstructured regions within the human CKM. Intrasubunit and intersubunit URPs identified between lysine residues are shown by red and blue lines, respectively. Functional and structural regions within individual CKM subunits are indicated. Intrinsically disordered regions (IDR) identified using IUPred3^66^ are shown based on propensity score, where residues with values greater than 0.5 are predicted to be disordered. Protein regions with more than 50 amino acids missing in the density maps are colored in white. **(B)** XL-MS map of the cMED- CKM, including MED12 and MED13, with all identified intersubunit and intrasubunit crosslinks. The color of the Mediator subunits is the same as that used in Figure 1C. **(C)** A close-up view of crosslinks occurring between H-lobe/MED13 and the cMED Hook. The Cα-atoms are shown by spheres. The residue numbers corresponding to crosslinked Lysines are shown as indicated. K1293 of MED13L corresponds to K1250 of MED13. **(D)** Intrasubunit (purple) crosslinks of MED13 and MED13L and their intersubunit crosslinks (green) with cMED subunits. The IDR regions of MED13 and MED13L are colored in gray. The color of the Mediator subunits is the same as that used in Figure 1C. **(E)** Crosslinks mapped between cMED and CKM are projected onto the cMED- CKM structure. Crosslinks occurring between cMED and MED13/13L IDRc are indicated by dashed lines. (**F**) Interaction site of Pol II with cMED overlaps with the location of URPs of MED13/13L IDRc. The location of Pol II on cMED is indicated by a transparent yellow surface.

Regarding the cMED-CKM (i.e. Mediator) complex, we identified 557 unique residue pairs (URPs), 86 of which were linked to either MED12L or MED13L; these URPs were not mapped onto the cMED-CKM structure, which does not include MED12L and MED13L (Figures 3B and S6E). Of the remaining 471 URPs, 277 were mapped onto the cMED-CKM structure with a low violation rate of 8.2% (Figure S6B and S6D). The remaining 194 unmapped URPs were found in either unstructured loops or large IDRs. Of the mapped 277 URPs, 214 and 56 occurred within cMED and CKM, respectively. Accordingly, only 7 URPs were mapped onto the Mediator structure between cMED and the CKM, four of which congregated between cMED Hook subunits and either the CKM H-lobe or MED13 (Figure 3C). These results support our structural model for human Mediator in which the main body of the CKM interacts with cMED through a limited contact area (Figure 1D).

Consistent with the localization of the H-lobe next to the cMED Hook, K762 and K772 in MED12 HEAT-III were crosslinked to K14 and K53 of MED10 and K95 of MED19, respectively, while K735 of MED12L (corresponding to K762 of MED12) was also crosslinked to K14 of MED10 (Figures 3C and S6E). Additionally, K1293 of the MED13L L2 domain (corresponding to K1250 of MED13) was crosslinked to K76 of MED14, which agrees with the localization of MED13 next to the Hook tip (Figure 3C). Significantly, both CDK8 and CCNC were extensively crosslinked to MED12 and MED13, but not to cMED subunits. These results are in line with our Mediator complex structure wherein the CKM binds to the cMED Hook in an orientation that results in displacement of the Kinase-lobe away from the main body of cMED (Figure 1D). Nevertheless, we cannot exclude the possibility that a small pool of the CKM may bind to cMED in a distinct manner based on some of the other identified crosslinks, i.e., one URP between CCNC and MED20, as well as three URPs between MED13/13L and MED14/17/24 (Figure S6B).

It is interesting to note that in the presence of cMED the IDRs, but not the Ago domains, in MED13/13L were extensively crosslinked to the Head and Middle module subunits in cMED (Figure 3D). By contrast, the majority of MED12 IDRs in the Mediator complex exhibited no crosslinks, suggesting flexible conformations without forming interactions with cMED. By mapping all of the URPs forged by the MED13/13L IDRs onto the cMED structure, we discovered that nearly all of these URPs are clustered at the central region of the Head module, with two URPs near the Knob of Middle module (Figure 3E). These findings suggest that the CKM allocates the MED13/MED13L IDRs to contact cMED at both its Head and Middle modules. Strikingly, the contacts formed between the MED13/13L IDRs and the Head module significantly overlap with the interaction site of Pol II on cMED, indicating their potential to influence Pol II binding (Figure 3F). This apparent binding mode contrasts with that observed in yeast where the CKM was crosslinked only to the cMED Middle module without overlapping the Pol II binding surface^36^. In addition, prior low-resolution structures of the yeast Mediator revealed a significant overlap between the main bodies of the CKM and Pol II^8,17^. Therefore, it is likely that human and yeast Mediator employ different mechanisms through which the CKM inhibits pol II binding.

### MED13 IDR binds cMED stably and competes with Pol II/MED26 for cMED binding

The ability of the MED13 IDR to interact with the Head and Middle modules in cMED could explain not only species-specific differences in CKM-mediated inhibition of Pol II binding, but further resolve an apparent discrepancy concerning the principal cMED interface within the CKM. Thus, while our structure identified MED12 as the major subunit within the CKM for cMED Hook binding, MED13 has nonetheless previously been reported to be the primary cMED interface based on subunit deletion and pull-down experiments^7,37,38^. To resolve these issues and establish whether and how MED13 and its IDR contributes to interactions between the CKM and cMED, we carried out a series of pull-down experiments combined with glycerol gradient centrifugation and mass spectrometry analysis. As shown in Figure 4A, recombinant full-length MED13 expressed in and purified from insect cells can pull-down cMED proteins from 293-F cell nuclear extract, suggesting its direct interaction with cMED. To identify the cMED-binding region (MBR) on MED13, we first expressed in and purified from insect cells the N-terminal (MED13-N; residues 1-1320) and C-terminal (MED13-C; residues 1070-2174) portions of MED13 that share an overlapping region of the L2 domain (Figure 4B). Pull-down experiments using these two MED13 domains and 293-F cell nuclear extract revealed that that MED13-N, but not MED13-C, bound cMED, thereby delimiting the MBR on MED13 to residues 1-1069 (Figure 4C left). To further delimit the MBR on MED13, we performed GST pull-down experiments using smaller MED13 fragments. This analysis revealed that MED13 residues 500-1069 corresponding to the C-terminal portion of its IDR (hereafter called IDRc) and two derivative fragments therein, residues 500-750 and 840-1069 (referred as MBR1 and MBR2, respectively), possess cMED binding capability (Figure 4C right). Flag-tagged IDRc, ectopically overexpressed in 293-F cells, was able to co-immunoprecipitate endogenous cMED, forming a stable complex that could be further purified through glycerol gradient centrifugation (Figure 4D). Additionally, the region in MED13L (residues 511-1107) corresponding to the MED13 IDRc also exhibited cMED interaction ability (Figure 4E). Collectively, these results show that the MED13 IDRc, through its constituent MBR1 and MBR2 domains therein, binds directly and stably to cMED.

**Figure 4.**
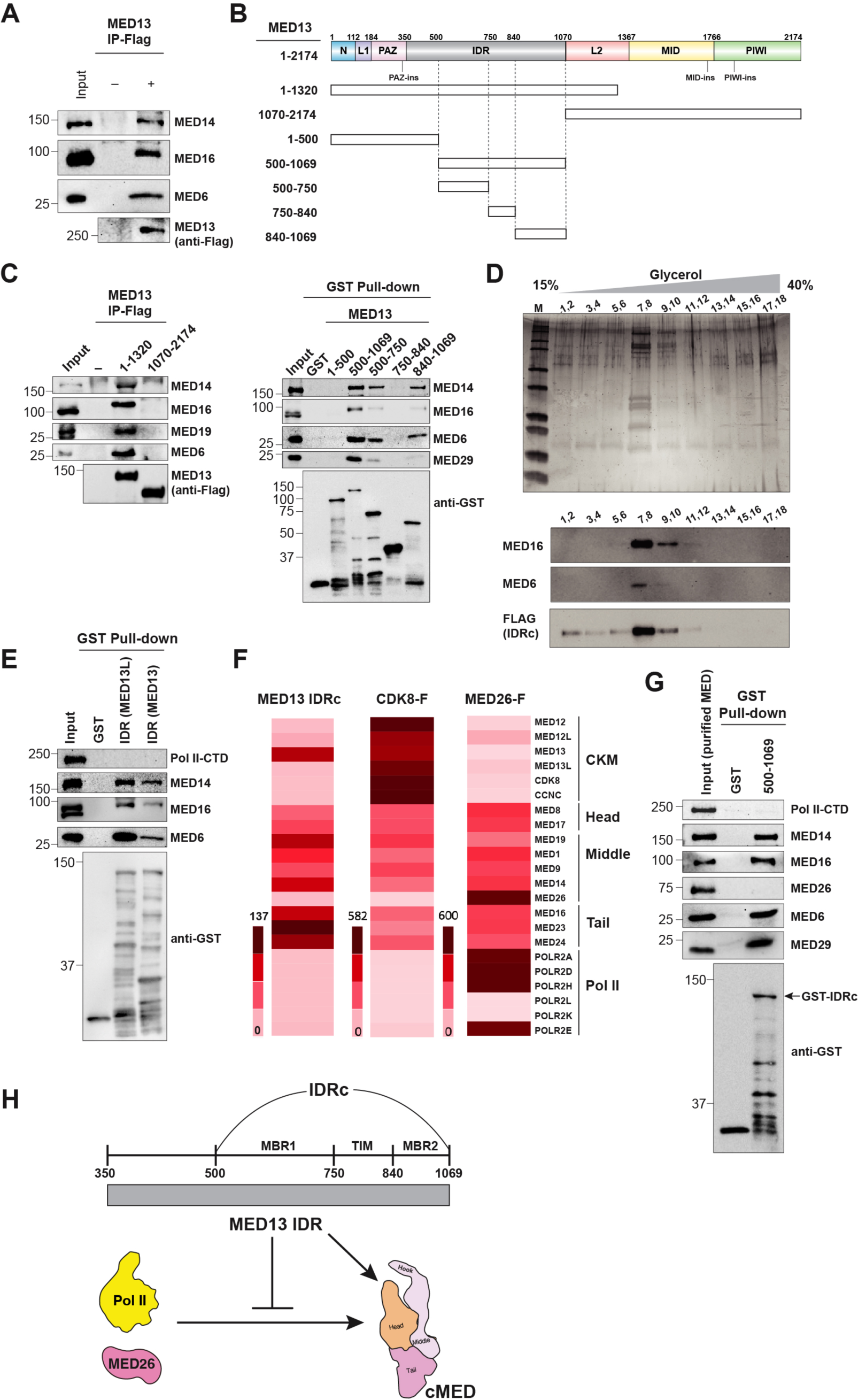
Interaction of cMED with the MED13 IDR precludes Pol II/MED26 binding. (A) Full-length MED13 interacts with cMED proteins. Purified recombinant Flag-tagged MED13 immobilized on anti-Flag M2 beads was used to pull-down cMED proteins from 293-F cell nuclear extract. **(B)** MED13 fragments used for cMED interaction analysis. **(C)** MED13 pull-down experiments. Left, the N-terminal portion (residues 1-1320) of MED13 associates with cMED as revealed by Flag IP analysis. Right, the IDRc, MBR1, and MBR2 of MED13 interact with cMED as shown by GST pull-downs. MED13 fragments were immobilized on anti-Flag M2 or Glutathione agarose beads to pull-down cMED proteins from the 293-F cell nuclear extract. **(D)** Interaction of the MED13 IDRc with cMED as revealed by glycerol gradient centrifugation. Gradient fractions were processed for SDS-PAGE and silver-staining (top) or immunoblot analysis with FLAG- and cMED subunit-specific antibodies as indicated (bottom). The distribution of Flag- tagged MED13 IDRc and cMED subunits are indicated. **(E)** GST pull-down experiment reveals cMED-binding ability of both MED13 and MED13L IDRc. The corresponding IDRc of MED13 or MED13L was immobilized on Glutathione agarose beads and used to pull-down cMED proteins present in 293-F cell nuclear extract. **(F)** Heatmap representing the scaled abundance of cMED, CKM, and Pol II proteins purified through Flag-tagged MED13 IDRc, CDK8 or MED26 followed by glycerol gradient centrifugation and mass spectrometry analysis. **(G)** MED13 IDRc-bound cMED precludes Pol II and MED26 association. GST-MED13 IDRc was immobilized on Glutathione agarose beads and used in pull-down assay with partially purified cMED comprising CKM-, as well as Pol II and MED26-bound cMED proteins. **(H)** A model showing capabilities of MED13 IDRc to interact with cMED and preclude Pol II/MED26 binding.

To fully characterize proteins that associate with the MED13 IDRc purified through anti-Flag immunoprecipitation and glycerol gradient centrifugation, the cMED-containing fractions were subjected to mass spectrometric analysis (Figure 4F left and Data S1). Intriguingly, IDRc- associated proteins included nearly all cMED subunits but neither MED26 nor Pol II, which is similar to the results obtained using Flag-tagged CDK8 (Figure 4F middle). In contrast, cMED proteins purified through Flag-tagged MED26 contained Pol II, but not CKM, albeit a small amount of MED13L was detectably present (Figure 4F right). This analysis indicates that the MED13 IDRc by itself, similar to the intact CKM, is able to compete with Pol II/MED26 for cMED binding. To confirm this, we conducted a pull-down assay using GST-IDRc and 293 cell-derived Flag-MED7-specific immunoprecipitates that include a mixture of CKM-, Pol II-, and MED26-bound cMED complexes. We found that indeed the MED13 IDRc binds only to Pol II- and MED26-free cMED (Figure 4G). Altogether, these results reveal a novel functional role for the MED13 IDRc in cMED binding and further confirm that that cMED interactions with Pol II/MED26 and the MED13 IDRc are mutually exclusive (Figure 4H).

### The CKM dispatches its MED13 IDR to occupy Pol II CTD and MED26 binding sites in cMED

To understand how human CKM inhibits the binding of Pol II to cMED, we first docked the CKM structure onto that of the human cMED-PIC (Figure 5A). Intriguingly, the main body of the CKM when bound to the cMED Hook is located >70Å away from Pol II and MED26 with no overlap (Figure 5A). This contradicts prior studies in yeast suggesting that steric hindrance by the main body of the CKM inhibits Pol II binding ^8,31,36^. Thus, it is likely that in humans the CKM precludes Pol II and MED26 association with cMED not through its main body, but rather through the cMED-IDRc, as revealed by our earlier protein binding analyses (Figure 4). This possibility is further supported by our XL-MS analysis of the cMED-CKM complex that revealed several URPs between the IDRc of MED13/13L and the Pol II/MED26 binding regions on cMED (Figure 3E and 3F). For example, within MBR1, K619 of MED13 and K659 and K678 of MED13L were linked to MED6, MED17, MED18, and MED22 around the Neck of the Head module where Rpb4/7 binds in the cMED-PIC structure (Figure 5B). Within MBR2, K932 and K947 of MED13 were crosslinked to MED31 and MED10, respectively, near the cMED MED21-MED7 Hinge domain where MED26 binds (Figure 5B). These results suggest that MBR1 and MBR2 of the MED13 IDRc are positioned around the Head and Middle modules, respectively, to preclude Pol II and MED26 binding.

**Figure 5.**
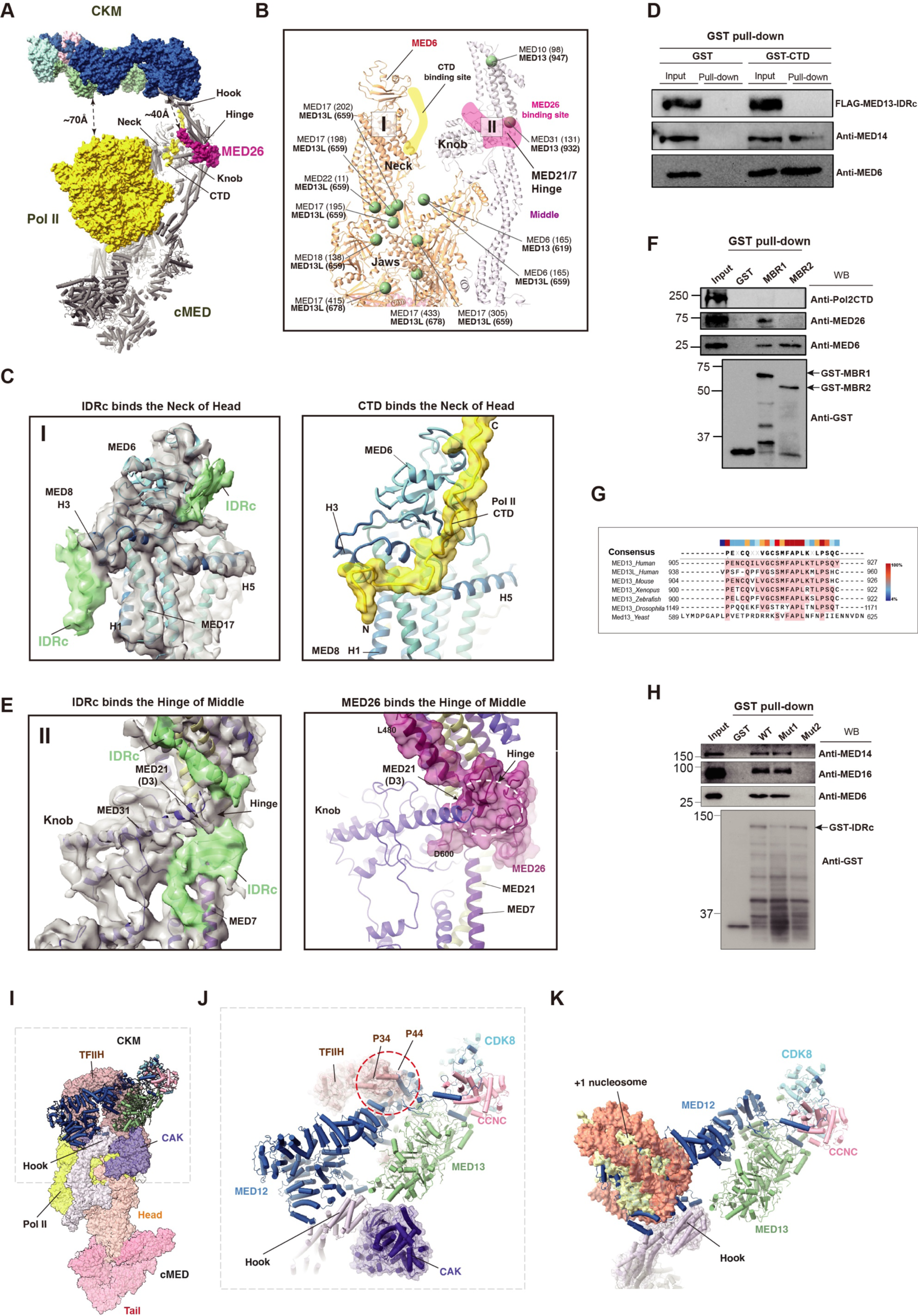
The MED13 IDR and CKM main body occupy Pol II/MED26 binding sites and affect PIC assembly, respectively. (A) Superimposition of the cMED-CKM and cMED-PIC [PDB 7ENC] structures by aligning their Hook domains reveals that the CKM main body does not overlap with Pol II (yellow) and MED26 (magenta). Distances between the CKM main body and either Pol II or MED26 are indicated. For clarity, cMED within cMED-CKM complex was omitted. **(B)** Crosslinks mapped between cMED and MED13/13L IDRc are projected onto the cMED-CKM structure. Specific lysine residues are indicated. The CTD (orange) and MED26 (purple) binding sites are indicated. **(C)** Left, a close-up view of the Neck domain of the Head module within cMED-CKM showing the specific location of extra densities (green maps) that correspond to the putative MED13*/*13L IDRc. Right, structure of the CTD (yellow) bound to the Head module within the cMED-PIC [PDB 7ENC]. **(D)** Pol II CTD-bound cMED precludes MED13 IDRc binding. GST-Pol II CTD was immobilized on Glutathione agarose beads and used in pull-down assay with nuclear extract from 293-F cells expressing Flag-tagged MED13 IDRc. **(E)** Left, close-up view of the Hinge domain of the Middle module within cMED-CKM shown with its corresponding density map (gray). The potential IDRc density is colored in green. The location of the MED21 D3K mutation is indicated. Right, structure of MED26 (magenta) bound to the Middle module Hinge domain within the cMED-PIC [PDB 7ENC]. Both locations of Knob and Hinge domains within cMED-CKM are indicated in (B). **(F)** GST pull-down experiment showing interaction of MED13 MBR1 and MBR2 with cMED. GST-MBR1 and GST-MBR2 were immobilized on Glutathione agarose beads and used to pull-down cMED proteins from 293-F cell nuclear extract. **(G)** Sequence alignment of conserved region within the MBR2 of the MED13 IDRc. **(H)** GST pull-down experiment reveals a conserved region within MBR2 is involved in cMED binding. WT or mutant MED13 IDRc was immobilized on Glutathione agarose beads and used to pull-down MED proteins present in 293-F cell nuclear extract. Mut1: F847D/S848Y/P849D mutant. Mut2: M916D/F917D/A918Ymutant. **(I)** Docking of human CKM onto the Hook of the human cMED- PIC [PDB 7LBM]. **(J)** The Central-lobe of the CKM partially clashes with the p34 and p44 subunits of TFIIH. **(K)** The H-lobe significantly overlaps with the proximal +1 nucleosome [PDB 8GXQ].

To determine the location of potential MED13 IDRc densities on cMED, we used focused 3D classification and refinement as well as fitting of the cMED structure into individually refined Head, Middle, and Tail maps (Figures S2F and S2G). Although a few URPs were detected between the Jaws of the Head module and the MED13L IDRc, no continuous extra densities were observed in this area (Figure 5B). By contrast, several extra densities were observed in the Neck domain of the Head module, as well as the MED21/7 Hinge next to the Knob of the Middle module, but not in the Tail module. More specifically, we identified two IDRc-like densities in the Head domain of cMED: an α-helix-like density located between H1 and H3 of MED8 near the central region of the Head, and a β-sheet-like density next to a conserved region on MED6 at the top of the Head (Figure 5C left). Notably, the helical density of the IDRc adjacent to MED8 agrees with several URPs between the MED13L IDRc and the bottom portion of the Neck domain as well as two additional URPs linking K619 of MED13 and K659 of MED13L to K165 of MED6 (Figure 5B). Since the Pol II CTD is located between the Head and Middle modules^26,39,40^, we investigated the possibility that the IDRc overlaps with the CTD by aligning structures of the cMED-PIC and the yeast Head-CTD complex with our model (Figures S7A and S7B). Strikingly, we found that the IDRc density adjacent to MED6 overlaps with the C-terminal region of the CTD (Figures 5C right and S7C). Furthermore, the IDRc density identified between H1 and H3 of MED8 occupies the same position where the N-terminus of the CTD binds in the structure of the yeast Head-CTD complex (Figures S7C). Since the cMED-Pol II interaction requires the CTD engaged between the Head Neck and Middle Knob following cMED structural rearrangements^26–28^, the IDRc bound to the Neck could preclude both CTD binding as well as CTD-induced cMED conformational changes. To further test this possibility, we performed an interaction analysis using GST-CTD to pull-down Mediator proteins from nuclear extract of 293-F cells in which the MED13 IDRc was overexpressed. This analysis revealed that the GST-CTD captured cMED proteins, but not the MED13 IDRc, (Figure 5D), supporting our structural findings that the MED13 IDRc obstructs the interaction of Pol II with cMED by occupying the CTD-binding site.

Beyond the Neck, we also identified additional densities at the N-terminus of MED21, which are located adjacent to the Knob of the Middle module (Figure 5E left). These densities include three β-strands that bind to the MED21/7 Hinge and two helices that contact the base of the Hook and the C-terminal region of MED7, respectively. The IDRc density at this position is substantiated by two URPs: K932 and K947 of MED13 crosslinked to K131 of MED31 and K98 of MED10, respectively (Figure 5B). It is noteworthy that these densities significantly overlap with the C-terminal domain of MED26, suggesting that the IDRc and MED26 compete for the same binding site on cMED (Figure 5E right). This structural observation not only explains why Mediator proteins purified through the MED13 IDRc did not contain MED26 (Figure 4F left), but also provides an explanation for the prior observation that a MED21 D3K mutation within the cMED Hinge disrupted both MED26 and MED13 binding^41^. We believe that these densities belong to the IDRc of MED13/13L because their localizations agree with our XL-MS analysis of cMED- CKM as describe above, and they were not shown in published maps of human cMED.

Our binding analysis has identified two cMED binding regions (MBR1 and MBR2) within the IDRc (Figure 4C). To further understand the roles of MBR1 and MBR2, we carried out Mediator interaction analyses and assessed their respective impact on Pol II and MED26 binding individually. Our findings show that MBR1 exclusively impacts Pol II binding without affecting MED26, whereas MBR2 can inhibit both Pol II and MED26 binding (Figure 5F). This result, along with our XL-MS analysis, suggests that MBR1 and MBR2 occupy the CTD binding site on the Head module and MED26 binding site at the Hinge domain, respectively. We also examined whether a highly conserved region (residues 915-920) in MBR2 near K932 of MED13 is involved in cMED interaction. To do this, we introduced a triple mutation (M916D/F917D/A918Y) within MBR2 of the IDRc and examined its impact on cMED binding (Figure 5G). Our results show that the IDRc mutant was significantly impaired for its interaction with cMED, indicating that this conserved region is involved in binding to the Hinge (Figure 5H). By contrast, an unrelated triple mutation (F847D/S848Y/P849D) located between MBR1 and MBR2 within the IDRc had no impact on the IDRc-cMED interaction, consistent with our prior binding assays indicating that the region between MBR1 and MBR2 does not contribute to cMED binding (Figure 4C right).

Altogether, our findings clearly explain the structural basis for mutual exclusivity among cMED interactions with the CKM and Pol II-CTD/MED26, highlighting a new and an important function for the MED13 IDRc as a key determinant of this differential binding.

### The CKM affects cMED-PIC assembly via steric hindrance with TFIIH and +1 nucleosome

To understand the potential impact of the CKM main body on cMED-PIC assembly, we assessed whether and how the CKM imposes any structural clashes with PIC components (Figure 5I). We found that the CKM can occupy a hollow area located between the main body of TFIIH and its CAK module, which contains CDK7, CycH, and MAT1^24,26,27^. Compared to CAK that contacts the edge of Hook, the CKM instead binds to the top and tip of the Hook without experiencing any structural clashes with CAK, suggesting that CKM binding does not affect the CAK-cMED interaction (Figure 5J). However, we found that MED12 in the Central-lobe of the CKM partially clashes with the p34 and p44 domains of TFIIH (Figure 5J). Thus, CKM binding is likely to affect cMED-PIC assembly through steric hindrance with TFIIH, an observation that could explain the abilities of MED12 to inhibit cMED-dependent transcription activity and influence chromatin occupancy of the cMED^7,42^. Additionally, we superimposed our cMED-CKM model onto the structure of the cMED-PIC in complex with +1 nucleosome^43^, located 40 bp downstream of the TSS, and found that the MED12 H-lobe significantly overlaps with the nucleosome (Figure 5K). Based on these structural considerations, it appears that the main body of the CKM needs to detach from the Hook prior to cMED-PIC formation due to the steric hindrance with TFIIH and the proximal +1 nucleosome.

### Human MED12 activates CDK8 without using a long activation helix

CDK8 and its paralog, CDK19, are the only enzymatic subunits present in the Mediator complex, and our structure revealed that CDK8 is located distally from the cMED and connected to the Hook through MED12 (Figure 6A). Within the Kinase-lobe of the CKM, MED12N adopts an extended conformation, which enables it to contact CDK8 near the T-loop through a coil followed by a short α-helix H1 (Figure 6B). Furthermore, MED12N makes numerous interactions with CCNC via a longer α-helix H2. Recently, the yeast CKM structure has revealed the molecular mechanism of Med12-dependent Cdk8 activation^31^. Herein, we show that human MED12N adopts a different conformation than that of yeast in order to bind and activate CDK8. Thus, unlike the N-terminal region of yeast Med12 that uses a long activation helix to stabilize the CDK8 T-loop, the N-terminal region of human MED12 instead adopts a coil structure followed by a short α-helix H1 to activate CDK8 (Figure 6C). To achieve this, the RHYT segment of human CDK8 undergoes a structural rearrangement induced by several contacts with the main chain of N40 and V41 of MED12 H1, as well as adjacent MED12 residues, D34, L36, and G44 (Figure 6D). These interactions alter the side chain orientation of Y211 in the RHYT segment, enabling a precise network of intramolecular interactions with the CDK8 T-loop. More specifically, Y211 in the RHYT segment directly contacts a conserved residue, R150, which in turn interacts with D191, a T-loop residue that is essential for kinase activation (Figure 6E). Notably, the tip of the CDK8 T-loop is directly contacted by MED12 through two residues, F45 and N47, which lie next to H1. Thus, the T-loop is well-defined in the electron density map (Figure S3D) and assumes an activated conformation in which residues 173-175, next to the ATP-binding site, adopt a “DMG-Asp-in” state, that is poised to coordinate Mg^2+^/ATP (Figures 6F). Next to the DMG motif, the flip of F176 toward the active site may participate in substrate binding. In contrast, the RHYT segment within the crystal structure of human CDK8/CCNC alone is configured in a different conformation that can potentially clash with the T-loop resulting in an inactive CDK8 (Figure S5B). Thus, we believe that human MED12N utilizes a coil to alter the conformation of the CDK8 RHYT segment, which, in turn, reshapes the active site by stabilizing the T-loop in an active conformation.

**Figure 6.**
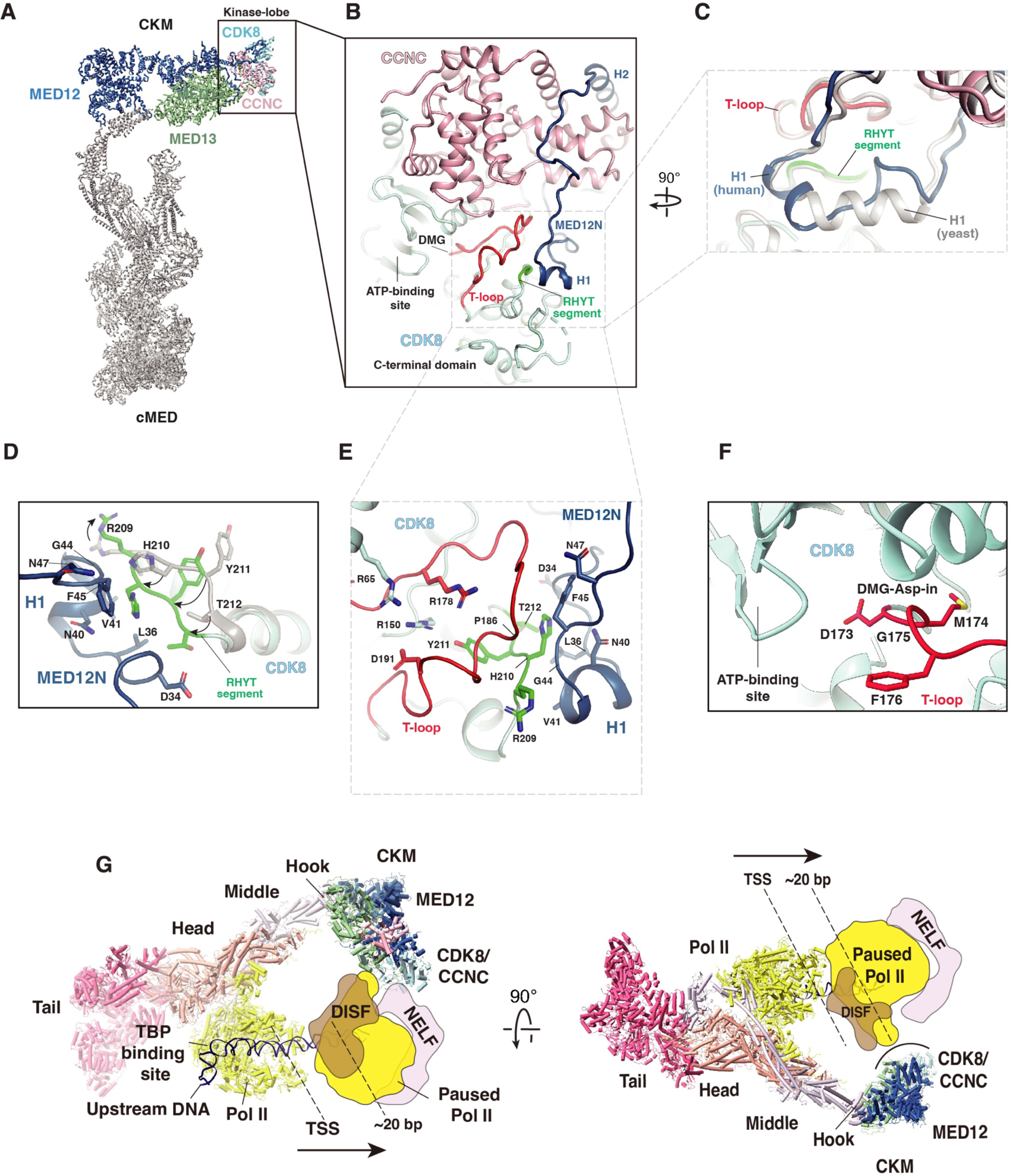
Activation mechanism of human CDK8 and its positioning downstream of the TSS. (A) Positional location of Kinase-lobe in the Mediator complex. **(B)** The structure of CDK8/CCNC bound by MED12N (blue). The CDK8 T-loop and RHYT segment are colored in red and green, respectively. **(C)** MED12N in humans adopts a conformation distinct from that in yeast for CDK8 activation. MED12N of yeast [PDB 7KPX] and humans are colored in white and blue, respectively. **(D)** The RHYT segment of CDK8 undergoes a structural rearrangement induced by MED12N. **(E)** A close-up view of the CDK8 T-loop and RHYT segment contacted by MED12N. Three CDK8 residues (R150, R178, Y211) important for coordinating an activated T-loop conformation are shown. MED12N residues involved in interactions with the CDK8 T-loop and RYHT segment are indicated. **(F)** The T-loop in a “DMG-Asp-in” conformation indicates active CDK8. **(G)** A putative model showing CKM and cMED as well as positions of Pol II (yellow) at the promoter and paused 20 bp downstream of the TSS. The structure of human cMED-PIC [PDB 7ENC] was used as a template in which the cMED was reverted to its free-form conformation. The putative TSS (+1) and 20 bp downstream of the TSS were indicated on the SCP promoter DNA^67^. The CKM was docked onto the cMED Hook, where it was found to be located approximately 30 bp downstream of the TSS. The structure of paused Pol II-DISF-NELF [PDB 6GML] was used to model the paused Pol II. Other PIC components were removed for clarity.

### The CKM in Mediator is positioned adjacent to promoter-proximal paused Pol II

Previous reports have indicated that cMED-CKM, but not cMED alone, is required for recruitment of pTEFb to alleviate Pol II pausing^18^. Additionally, CDK8 can phosphorylate multiple pausing-related factors, such as NELFA and NELFB^22,23^. To gain structural insight into CKM-mediated gene activation through regulation of Pol II pausing and release, we first docked the CKM onto the Hook of the cMED-PIC structure^26^. In this structure, a Pol II molecule bound by NELF was modeled and located ∼20 bp downstream of the TSS ^26^. Interestingly, we found that the main body of the CKM on the cMED Hook is localized ∼30 bp downstream of the TSS, where it is suitably positioned to phosphorylate proteins such as NELF bound to paused Pol II (Figure 6G). Additionally, the presence of the CKM adjacent to the Pol II pausing site could contribute to recruitment of pTEFb. These structural observations provide molecular insight into the positive function of CKM and CDK8 in the regulation of Pol II pausing and release.

## DISCUSSION

The eukaryotic Mediator plays a central role in regulating Pol II-dependent gene transcription through its large cMED and dissociable CKM, which possesses a non-canonical CDK8 kinase. The cMED activates basal transcription by enlisting Pol II via a CTD-dependent interaction, promoting PIC formation, and stimulating CTD phosphorylation by TFIIH^3,44^. Recent structural studies have provided detailed insights into how the cMED and PIC are assembled to activate transcription^45^. Nevertheless, the mechanisms by which the CKM can both repress cMED activities prior to transcription initiation and positively influence gene expression, potentially in post-transcription events, remain unclear. In this study, we used single particle cryo-EM analysis to determine the structures of the human Mediator and its dissociable CKM. Our results provide a structural and molecular mechanistic basis to explain how the CKM interacts with cMED, represses cMED-dependent transcription, and supports MED12-dependent CDK8 activation.

The CKM, which reversibly associates with cMED, is present and conserved in all eukaryotes, yet it was originally thought to function solely as a transcriptional repressor^7,46^. Despite lack of definitive high-resolution structural information, steric hindrance involving occlusion of the Pol II binding site on cMED by the CKM main body has been considered the principal mechanism for CKM-mediated transcription repression^8,17,31,36^. In our human Mediator structure, the CKM binds to the Hook of cMED (Figure 1D). Intriguingly, the main body of CKM in this binding mode does not overlap with Pol II, and CKM binding does not induce any apparent cMED conformational changes. Accordingly, neither steric hindrance nor conformational changes induced by binding of the CKM main body can explain its inhibitory effect on cMED-dependent transcription. Instead, we find that the MED13 IDRc, through its bivalent association with cMED, directly occludes Pol II CTD and MED26 binding, thus revealing a hitherto unknown role for the MED13 IDRc as a modulator of cMED activity. Prior studies showing that MED26 is present in cMED, but not in cMED-CKM, along with those demonstrating that Pol II is enriched in cMED purified from cells expressing FLAG-MED26 suggest that MED26 may help to recruit Pol II to cMED^6,47^. Furthermore, MED26 has been implicated in TFIID and/or SEC (Super elongation complex) recruitment^11^ and it has also been shown to be essential for cMED-dependent transcription *in vitro* ^48^. Therefore, by blocking the binding of MED26 to cMED, the MED13 IDRc may, in addition to inhibition of Pol II recruitment, impose an additional layer of regulation upon cMED-dependent gene expression.

Since the IDR is present in all MED13 family proteins^49^, its function in precluding Pol II binding to cMED is likely to be conserved. Accordingly, dissociation of the MED13 IDR from cMED should be a critical regulatory step prior to Pol II binding or cMED-PIC formation in the transcription initiation process (Figure 7). Post-translational modifications (PTMs), including MED13 ubiquitination and/or phosphorylation as previously reported, represent possible ways to remove the MED13 IDRc from cMED^36,38^. Further studies on PTMs to the MED13 IDR and their possible influence on its association with cMED could reveal important mechanistic insight regarding the basis by which the MED13 IDR dissociates from cMED. The IDRc of MED13 contains two noncontiguous cMED-binding regions, MBR1 and MBR2, separated by a region (residues 750-840) with no apparent cMED-binding activity (Figure 4C right). Interestingly, this fragment was recently reported to contain a TND-interaction motif (TIM) for elongin A binding^50^, suggesting that MED13 IDRc may also exert positive regulation. On the other hand, despite a moderate sequence identity of ∼41% (Figure S8A) between the two corresponding IDRs of MED13 and MED13L, cMED proteins purified through MED26 revealed the presence of MED13L, but not MED13 (Figure 4F). This observation suggests that the functions of both IDRs are not wholly conserved, which broadly increases the capability and functionality of the CKM in gene regulation.

**Figure 7.**
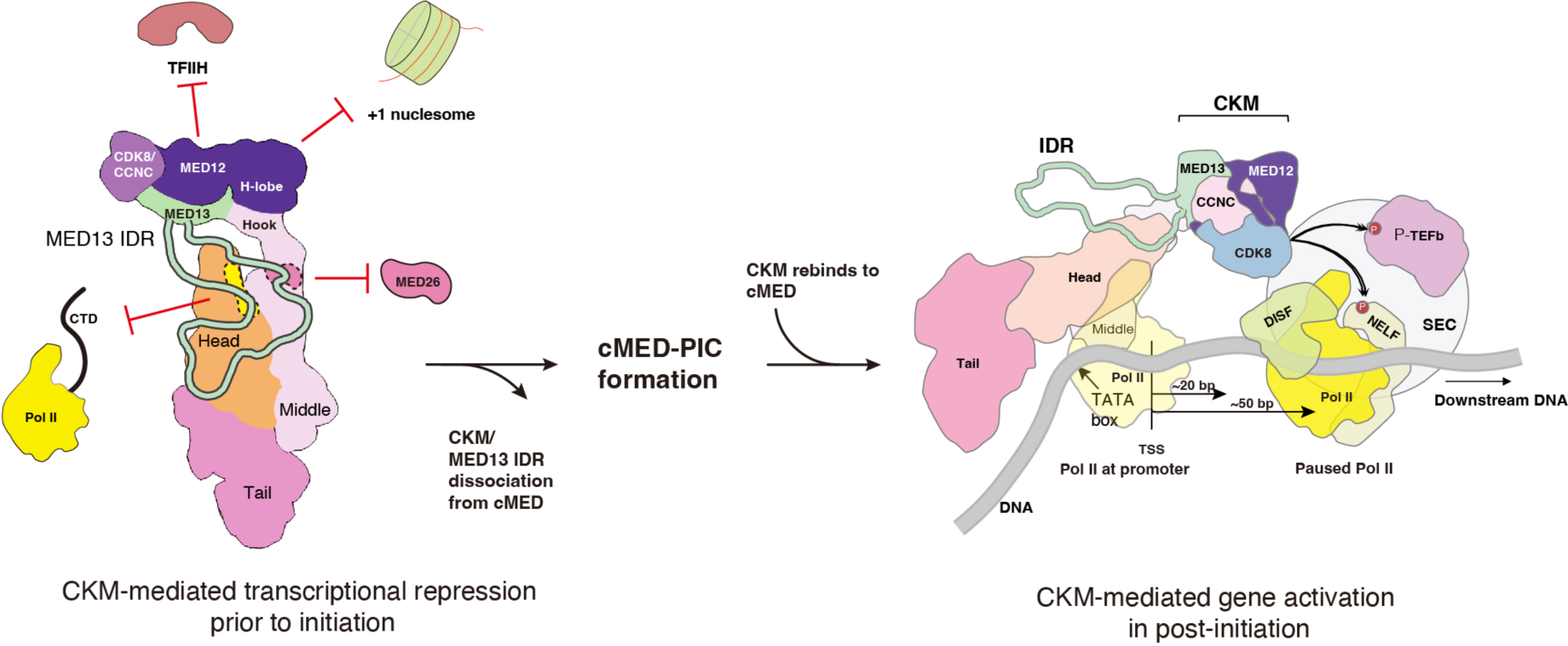
Models for CKM-mediated transcription repression and gene activation. Left, prior to transcription initiation, the CKM dispatches its MED13 IDR to occlude binding of both Pol II CTD and MED26 to cMED, thus inhibiting cMED-PIC formation. Both CTD and MED26 binding sites on cMED are highlighted by yellow and magenta dashed shapes, respectively. Dissociation of the MED13 IDR is required for cMED to bind Pol II and MED26 leading to cMED- PIC formation. Right, following promoter escape, Pol II pauses 20-50 bp downstream of the TSS. Interaction between the MED12 H-lobe and cMED Hook deploys the CKM downstream and proximal to the TSS where it is positioned to recruit the SEC and/or promote CDK8-dependent phosphorylation of pause-release-related factors, such as NELF proteins. DNA is shown by a gray line. For clarity, other PIC components are removed.

In yeast, cMED and the CKM are localized predominantly at upstream activating sequences which are bound by gene-specific activator proteins^51–54^. By contrast, in mammalian cells, both cMED and the CKM are stably associated with promoters, as indicated by ChIP-seq studies^53,55^. This finding is surprising because the CKM was previously believed to prevent the interaction of Pol II with cMED, resulting in repressed transcription. Furthermore, our cMED-PIC- CKM model indicates that the main body of the CKM needs to detach from the cMED Hook during PIC formation due to a minor steric clash with TFIIH (Figure 5J). Thus, it is likely in mammalian cells that the CKM may reassociate with promoters through its interaction with cMED after promoter escape whereupon it participates in post-initiation processes. This may explain why CKM association with promoters was observed in ChIP-seq studies.

In yeast, the CKM has been found to preferentially associate with highly transcribed genes^56,57^. In Arabidopsis, Med12 and Med13 act as positive gene regulators and are highly enriched in gene bodies near TSS^58^. Additionally, the CKM and its resident CDK8 participate in post-initiation events, such as Pol II pausing and release^20,21^. In this regard, CDK8 phosphorylates NELFA and NELFB, and inhibition of CDK8 activity was shown to increase Pol II pausing^22,23^. Furthermore, cMED-CKM was found to be required for recruitment of the SEC to alleviate Pol II pausing in response to a variety of stimuli^18^. A noteworthy observation from our structural studies is that the CKM is located downstream of the TSS and in close proximity to the +30 bp site where Pol II is commonly paused (Figure 7). This could facilitate CDK8 access to its phosphorylation its substrates, such as NELF factors bound to paused Pol II. The large surface of the CKM could also contribute to recruitment of the SEC. Since our Mediator structure shows that the CKM deploys both its MED12 H-lobe and MED13 IDRc to bind cMED at two distinct points of contact, it is possible that the main body of the CKM, via the MED12 H-lobe, could remain associated with the Hook even when the MED13 IDRc is dissociated from cMED. This would provide Pol II access to cMED for re-initiation and, in turn, allow the CKM to alleviate Pol II pausing. This structural observation could account for the positive function of the CKM in gene expression. Further studies will be required to determine the significance of the interaction between MED12 and Hook in relation to transcriptional pause and release.

Unexpectedly, we find that human MED12 and MED13 exhibit significant structural differences when compared to their corresponding subunits in the yeast CKM, probably because of their low sequence identities (12.7 and 13.0 % for MED12 and MED13, respectively). For example, two Zn-binding sites absent in the yeast CKM were identified in human MED12 and MED13. Furthermore, the overall structure of human MED13, including the structure of the L2 domain, is markedly distinct from that of its yeast counterpart. In this regard, whereas the L2 domain in yeast Med13 adopts a coiled structure resulting in a more compact overall architecture, the corresponding region in human MED13 consists of several α-helices that occupy and widen the central channel (Figure S5A). Additional studies will be required to establish the functional significance of the unique Ago-like structure of MED13. Finally, a comparative analysis of MED12 in the human and yeast CKM structures highlights several notable differences (Figure S5D). First, in contrast to yeast, the N-terminal region of MED12 in humans does not employ a long α-helix to activate CDK8. Instead, human MED12N primarily uses a coil followed by a short α-helix (H1) to bind the CDK8 RHYT segment; the corresponding region in the yeast is structured into a longer α-helix (Figure 6A). Second, unlike yeast Med12N, which engages in substantial interactions with Cdk8, the shorter human MED12N exclusively binds to the T-loop and RHYT segment (Figure S5D). Interestingly, these additional Cdk8 interactions forged by the longer Med12N in yeast are effectively replaced in the human CKM by an extended loop (residues 1616- 1630) from the MED13 PIWI domain (Figure S5D left), likely preserving additional CDK8 contacts required for complex stability. Finally, in contrast to the curved architecture characteristic of the yeast CKM H-lobe, the corresponding H-lobe of the human CKM adopts a more extended conformation with additional loops that likely mediate protein-protein interactions (Figure S5D right).

Due to sequence similarities in MED12/12L and MED13/13L, we were unable to distinguish MED12 and MED13 from their corresponding paralogs based on densities in the cMED-CKM. However, we noticed that the ordered regions within MED12 and MED13 exhibit high sequence identity with their corresponding paralogs (Figures S8B and S8D). This is also supported by AlphaFold models of MED12L and MED13L, which can be well-superimposed onto the cryo-EM structures of MED12 and MED13, respectively, except for the N- and C-terminal regions of MED12/12L and the IDRs of MED13/13L. (Figures S8C and S8E). Accordingly, the overall structures, cMED-binding modes, and CDK8 activation determined for MED12 and MED13 are likely similar to those of their corresponding paralogs.

The CKM in humans has been extensively linked to diseases, including cancer^59^. Our human CKM structure not only confirms the importance of the MED12 N-terminus in both binding and activation of CDK8, but also provides a structural framework to score the effects of disease-causing mutations on CKM function (Figure S9)^59^. Uterine fibroid hotspot mutations in MED12 ^60^ are highly clustered within or adjacent to H1 and critically interface with the CDK8 RHYT/T-loop region, providing a structural basis to explain the deleterious impact of these mutations on MED12-dependent CDK8 activation^14,61,62^. In our CKM structure, MED12 G44 that contacts the RHYT segment is next to the T-loop (Figure 6E). Accordingly, G44 mutations are likely to alter the T-loop and RHYT segment through their side chains, leading to impaired kinase activity, which explains why G44 is the most frequently mutated residue in uterine fibroids (Figure S9B)^14,59^. On the other hand, two human MED12 disease mutations (G958E and R961W) causative for the intellectual disability-multiple congenital anomaly disorder FG syndrome that disrupt interaction between cMED and activating non-coding RNAs are located within the H-lobe near the HEAT III-IV interface, implicating the H-lobe in functional RNA association (Figure S9A) ^63^. Additionally, one MED12 disease mutation H1729N causative for the blepharophimosis-intellectual disability disorder Ohdo syndrome could impair CKM stability by influencing Zn binding because the H1729 is involved in coordinating one of the MED13 Zinc-binding sites (Figure 2C)^33^.

CDK8 is a potent colorectal oncogene, spurring efforts to develop clinical inhibitors, such as type-II inhibitors that bind to the deep pocket adjacent to the ATP-binding site^64,65^. Recently, it has been reported that type-II inhibitors exhibit reduced efficacy toward the CDK8/CCNC when bound by MED12N^15^, but the reason for this phenomenon is unknown. To obtain a structural explanation, we fitted type-II inhibitors into the structure of human CKM and found that inhibitors bound at the deep pocket exhibit steric clash with residue M174 of the DMG motif (Figure S5C). Thus, MED12 binding stabilizes the active site in a “DMG-Asp-in” conformation which creates steric hindrance to type-II inhibitors, explaining the incompatibility of type II inhibitors for MED12-bound CDK8.

In summary, the human CKM can enhance and repress cMED-dependent transcription through its multifunctional regulatory properties that include MED13 IDR-mediated control of cMED-Pol II/MED26 interactions and MED12-dependent CDK8 kinase activity. Prior to initiation, CKM-bound cMED is transcriptionally inhibited by the MED13 IDRc that occludes Pol II and MED26 association, thereby preventing cMED-PIC formation and recruitment of elongation factors. Accordingly, dissociation of the MED13 IDR from cMED likely represents a critical regulatory step in cMED-dependent transcriptional activation. In post-transcription initiation events, the CKM bound at the cMED Hook is positioned near the Pol II pausing site, providing a structural basis to explain CKM-mediated gene activation through recruitment of the SEC and CDK8-dependent phosphorylation of pausing-related factors.

## Acknowledgments

We thank the Structural Biology Imaging Center of the UTHealth McGovern Medical School for help with cryo-EM data collection. We thank Mass spectrometry facilities at the University of Pennsylvania, the University of Texas Health Science Center, and Baylor College of Medicine for conducting XL-MS analysis and protein identification. This work was supported by the Cancer Prevention Research Institute of Texas, Grant number 13127 to CPRIT Scholar in Cancer Research, Kuang-Lei Tsai, the Welch Foundation (AU-2050-20200401), and US National Institutes of Health grants R01 GM143587 (K-L.T.), HD087417 and HD094378 (T.G.B), and R01 GM123233 (K.M.), and NIH NRSA F31AG069390 (H.J.K.).

## Author Contributions

S.F.C. prepared the human CKM specimen for structural determination with help from H.C.T. and S.K.; T.C.C. prepared the human cMED-CKM specimen for structural determination with help from H.C.T. and S.K.; S.F.C., T.C.C., and K.L.T. performed all experiments related to high-resolution cryo-EM structural analysis, including cryo-EM grids preparation, data collection and processing, and model building and refinement. H.J.K., and K.M. performed XL-MS and analyzed the data; S.F.C., T.C.C., and H.C.T. designed and performed binding assay and kinase activity measurement; K.M., T.G.B., and K.L.T supervised the project; All authors discussed and interpreted results; S.F.C., T.C.C., H.J.K., K.M., T.G.B., and K.L.T. wrote the manuscript.

## STAR METHODS

### Lead Contact

Further information and requests for reagents should be directed to and will be fulfilled by the Lead Contact, Kuang-Lei Tsai.

### Materials Availability

Plasmids and strains generated in this study are available upon request from Lead Contact with a completed Materials Transfer Agreement.

### Data and Code Availability

- **Cryo-EM:** Cryo-EM maps of the human cMED-CKM, cMED-HookΔ, CKM-Hook, Head- IDR, and Middle-IDR were deposited to the EMDataBank with accession numbers EMD-xxxxx, EMD-xxxxx, EMD-xxxxx, and EMD-xxxxx, respectively. The cMED-CKM model built based on these maps were deposited to the RCSB Protein Data Bank with an accession number XXXX. Cryo-EM maps of the human CKM, Kinase-lobe, Kinase/Central-lobes, and H-lobe were deposited to the EMDataBank with accession numbers EMD-xxxx, EMD-xxxx, EMD-xxxx, and EMD-xxxxx, respectively. The CKM model built based on these maps was deposited to the RCSB Protein Data Bank with an accession number XXXX.
- **Crosslinking mass-spectrometry:** Raw data and analyses results were deposited to XXX.

## METHOD DETAILS

### Cell culture

The pcDNA3.1 plasmids encoding human Flag-tagged CDK8, MED13 IDR (residue 500-1069), MED7, and MED26 were generated and then transfected into Freestyle 293-F cells (HEK 293-F, ThermoFisher) using Lipofectamine 3000 (Invitrogen). Transfected cells were selected by treating 100 ug/ml G418 and maintained in FreeStyle^TM^ 293 expression medium with 50 ug/ml G418. All 293-F cells were maintained in shaking incubators at 37°C, humid atmosphere and 8% CO_2_ in a serum-free culture medium.

### Purification of Human cMED-CKM

10 Liters of Freestyle 293 cells expressing CDK8-Flag were harvested, washed with cold 1x PBS twice, and resuspended in 250 ml hypotonic buffer (10 mM Hepes pH7.9, 1.5mM MgCl_2_, 10 mM KCl, 0.5 mM Dithiothreitol, and protease inhibitors). After incubation on ice for 10 mins, the cells were homogenized in a Dounce homogenizer with 20 gentle strokes of the loose pestle. The cell lysate was centrifuged at 20,000 rpm for 30 mins at 4°C using a Beckman type 45Ti rotor, and the resulting pellet was washed by the hypotonic buffer twice, followed by centrifugation at 20,000 rpm for 10 mins at 4°C. Nuclear extract was prepared by following a protocol as described^68^. In brief, the pellet was gently suspended in 50 ml extraction buffer (25 mM Hepes pH7.9, 1.5mM MgCl_2_, 0.3 M KCl, 0.5 mM DTT, 0.1 mM EDTA, 0.01% NP-40, 10% glycerol, and protease inhibitors) and rotated end-over-end for 1 hr at 4°C. The nuclear extract was centrifuged 33,000 rpm for 1 hr at 4°C in a Beckman type 45Ti rotor. The supernatant was collected and incubated with 5 ml anti-Flag M2 beads at 4°C for 4 hours with end-over-end rotation. The beads were loaded into a column and washed 4 times with 30 ml of washing buffer (25 mM Hepes pH7.6, 1.5mM MgCl_2_, 150 mM NaCl, 10% glycerol, 0.2 mM EDTA, 0.01% NP-40, and 2 mM beta-mercaptoethanol). The cMED-CKM complex was eluted with elution buffer (25 mM Hepes pH7.6,

1.5mM MgCl_2_, 150 mM NaCl, 10% glycerol, 0.2 mM EDTA, 0.01% NP-40, 2 mM beta-mercaptoethanol, and 200 ug/ml 1x FLAG peptide) at 4°C for 1 hr with end-over-end rotation. cMED-CKM eluted from anti-Flag M2 beads was concentrated and then loaded on a 15%-40% glycerol gradient [20 mM Hepes pH7.6, 1 mM MgCl_2_, 300 mM NaCl, 0.25 mM EDTA, and 1 mM beta-mercaptoethanol, and 15%-40% (v/v) glycerol], and centrifuged for 17 hours at 30,000 rpm using a Beckman SW60 Ti rotor. The fractions were examined by SDS-PAGE and wester blotting. The peak fractions containing cMED-CKM complex were combined and dialyzed into buffer (20 mM Hepes pH7.6, 1 mM MgCl_2_, 300 mM NaCl, 0.25 mM EDTA, and 2.5 mM beta-mercaptoethanol). After dialysis, the cMED-CKM protein was examined by negative-stain EM, concentrated to ∼1 mg/ml, and subjected to cryo-EM specimen preparation.

### Expression and purification of recombinant human CKM

pFastBac1 plasmids encoding human CDK8-Flag, CCNC-6xHis, MED12-HA, and CBP-MED13 for baculovirus-based protein expression were generated and then transformed into DH10Bac competent cells (ThermoFisher) as previously described^69^. The bacmid DNAs isolated from white colonies were verified by DNA sequencing and used for transfection of Sf9 insect cells. After three rounds of viral amplification, four high-titer recombinant baculoviruses were used for co-infection of High Five cells (ThermoFisher). After 48 hours infection, 3 Liters of High Five cells were harvested and lysed with buffer A (20 mM Hepes pH 7.9, 150 mM NaCl, 0.01% NP-40, 0.1mM EDTA, 10% glycerol, and protease inhibitors) using dounce homogenizer. Lysates were clarified by high-speed centrifugation (sorvall ss-34 rotor) at 18,000 rpm for 1 hour. The supernatant containing recombinant CKM proteins was incubated with anti-Flag M2 affinity beads for 1 hour at 4°C in buffer A. The CKM was washed four times, eluted with elution buffer (buffer A containing 200 ug/ml 1x FLAG peptide), and analyzed by SDS-PAGE. The CKM eluate was subjected to 15%-35% glycerol gradient centrifugation in buffer [20 mM Hepes pH 7.9, 150 mM NaCl, 0.1mM EDTA, and 15%-35% (v/v) glycerol] with a SW60Ti rotor at 44,000 rpm for 16 hours at 4°C. The peak fractions containing CKM four subunits were pooled and concentrated, followed by buffer exchange into a buffer containing 20 mM Hepes pH 7.9, 150 mM NaCl, and 0.1mM EDTA for subsequent kinase assay and structure determination.

### Expression and purification of recombinant MED13 proteins

DNA fragments encoding residues 1-500, 500-1069, 500-750, 750-840, 1-1320, 1070-2174, and 840-1069, as well as two IDRc mutants (M916D/F917D/A918Y and F847D/S848Y/P849D) of human MED13 were generated by PCR amplification using full-length MED13 DNA as a template. Except for the residues 1-1320 and 1070-2174 of MED13 that were constructed into pFastBac1, the remaining MED13 fragments were ligated into pGEX6P-1 for expression in *E.coli*. All constructs were verified by DNA sequencing. For GST pull-down assays, each of the MED13 fragments, fused to the C terminus of GST protein, was expressed in *E. coli* BL21(DE3) followed by the procedure as described below.

### GST Pull-down assay

pGEX6P-1 plasmids encoding human CTD domain of Pol II RPB1 (residues 1593-1970) or MED13 fragments (residues 1-500, 500-1069, 500-750, 750-840, and 840-1069) were transformed into *E. coli* BL21(DE3) and Rosetta(DE3) pLysS cells, respectively. Protein expression was induced by addition of IPTG at a final concentration of 0.1 mM at 25°C for 3 hours. Cells were harvested and lysed by sonication. After centrifugation, the clarified lysate was incubated with 25 ul glutathione sepharose 4B beads (GE Healthcare) for 30 min at 4°C. The beads were washed three times using binding buffer (20 mM Hepes pH 7.9, 300 mM NaCl, 0.01% NP-40, 10% glycerol, and protease inhibitors) followed by incubation at 4°C for 3 hours with 400 ul 293-F nuclear extracts or partially purified Mediator proteins that contain CKM- or Pol II-bound cMED, which were obtained using MED7-Flag as described above. After binding, the beads were washed three times using binding buffer and then eluted using elution buffer containing 10 mM glutathione. The eluates were analyzed by SDS-PAGE and Western blotting using anti-GST antibodies (GenScript, A0086640), anti-MED14 antibodies (Bethyl Laboratories, A301-044A), anti-MED16 antibodies (Bethyl Laboratories, A303-668A), anti-MED6 antibodies (Santa Cruz, sc-390474), anti-MED26 antibodies (Cell Signaling, 13641S), and anti-Pol II CTD antibodies (8WG16).

### FLAG immunoprecipitation

High Five cells for expressing full-length, or N-terminal (residues 1-1320) or C-terminal (residues 1070-2174) regions of Flag-tagged MED13 were prepared, harvest, and lysed following the same procedure as described above. After centrifugation, each supernatant was collected and incubated with 30 ul FLAG-M2 beads for 30 minutes at 4°C. After immobilization, the Flag beads were washed three times with binding buffer B (25 mM Hepes pH7.6, 1.5mM MgCl_2_, 300 mM NaCl, 10% glycerol, 0.2 mM EDTA, 0.01% NP-40, 2 mM beta-mercaptoethanol) and then incubated with the 293-F nuclear extracts for 3 hours at 4°C. The beads were washed three times with binding buffer B and then eluted with the same buffer containing 200 ug/ml 1x FLAG peptide. The eluates were analyzed by SDS-PAGE and Western blotting using anti-FLAG M2 antibodies (Sigma-Aldrich, F1804), anti-MED14 antibodies (Bethyl Laboratories, A301-044A), anti-MED16 antibodies (Bethyl Laboratories, A303-668A), and anti-MED6 antibodies (Santa Cruz, sc- 390474).

### Negative stain EM

3 μl of diluted cMED-CKM protein sample (∼0.03 mg/mL) in buffer (20 mM Hepes pH7.6, 200 mM NaCl, and 0.1 mM EDTA) was applied onto a carbon-coated and glow-discharged 300 mesh Cu/Rh grids (Ted Pella). After 1 min waiting, the excess of the sample was blotted using a piece of filter paper. Immediately after blotting, 3 μl of 2% uranyl acetate was applied onto the grid and then incubated for 1 min. The same blotting and staining steps were repeated two times. Then the grid was blotted dry from the edge with filter paper and allowed to dry. The stained grid was imaged on a JEOL 1400 electron microscope with a Rio9 camera (Gatan).

### Cryo-EM specimen preparation, data collection and image analysis

In brief, 3.0 μl of human CKM (∼1 mg/ml) in buffer (25 mM Hepes pH 7.4, 200 mM NaCl, and 0.1 mM EDTA) were directly applied onto freshly grow discharged 300 mesh C-flat^TM^ holey carbon grids (CF-2/1-3C). Grids were blotted for 3–4 s at 4°C with 100% humidity and flash-frozen in liquid ethane using a Vitrobot mark IV (FEI). For cMED-CKM, 3.0 μl of concentrated protein (∼0.5 mg/ml) in buffer (25 mM Hepes pH 7.6, 150 mM NaCl, 0.1 mM EDTA, 2 mM MgCl_2_, and 1 mM beta-mercaptoethanol) were applied onto grow discharged 300 mesh C-flat holey carbon grids. Frozen grids of cMED-CKM were prepared following the same procedure as described for the CKM above.

For cryo-EM data collection of CKM, the grids were imaged on a 300 kV Titan Krios electron microscope (FEI) using a GIF Quantum K2 direct electron detector (Gatan) operating in counting mode. Images were automatically collected at 0.6–3.0 um underfocus values with a nominal pixel size of 1.07 Å per pixel using EPU (Thermofisher). Each image was exposed for 8 s with a total dose of approximately 64 electrons per Å^2^, which was fractioned into 40 frames. MotionCor2 was used to align frames^70^. The parameters of contrast transfer function (CTF) for each image were estimated using the program Gctf^71^. Initial particle picking by crYOLO^72^ resulted in ∼1.3 million particles. 2D clustering in cryoSPARC^73^ was carried out to obtain a stack of ∼30K images that was used to generate initial 3D models of CKM. 2D and 3D classification were carried out in RELION^74^, and identified a set of 107,310 images that were run through 3D refinement, Bayesian polishing, and CTF refinement to obtain a 3D map of overall CKM at an average resolution of 3.8 Å, in which the secondary structures of the H-lobe were poorly defined due to mobility and conformational heterogeneity. For Kinase- and Kinase/Central-lobes of the CKM, the 107,310 images were further 3D classified and refined with a focused mask that covered corresponding regions, resulting in final maps at 3.8 and 3.6 Å resolution, respectively. For the H- lobe of CKM, we also started from the 107,310 images and performed 3D classification with a mask that only covered the H-lobe using RELION^74^. A final stack of 68,316 images were selected to run 3D refinement with the same mask, resulting in a final H-lobe map at 6.5 Å resolution. The resolutions of final 3D maps were estimated using gold standard Fourier Shell correlation (FSC) curves with 0.143 criteria^75^. RELION or Cryosparc were used to calculate local resolutions.

For cryo-EM data collection of the cMED-CKM, the grids were imaged on the same Titan Krios electron microscope equipped with a K2 direct electron detector with the following settings. Images were collected at 0.6-2.8 um underfocus values with a nominal pixel size of 1.42 Å per pixel using EPU counting mode. Each image was exposed for 12 s and fractioned into 48 frames with a total dose of approximately 52 electrons per Å^2^. The movie frames were aligned by MotionCor2^70^, resulted in a total of 15,822 micrographs. crYOLO^72^ was used for particle picking that resulted in a total of 467,838 particles. Multiple rounds of 2D classification in RELION/Cryosparc were performed to remove particles that lacked clear features, resulting in a stack of 290,607 particles. A subset of 25,316 particles was used for 3D initial model generation using Cryosparc. An initial model showing the CKM bound to cMED was low-pass filtered to 60 Å and used as a reference for carrying out 3D classification in RELION. This resulted in five classes with much better structural features than the others. These classes were combined into a stack of 105,401 images, which were subjected to 3D refinement that resulted in a 6.4 Å reconstruction. CTF refinement and Bayesian polishing were carried out to generate 105,401 shiny particles. 3D refinement of these shiny particles resulted in a 4.8 Å reconstruction, in which the CKM density was poor due to flexibility. A mask that masked out the CKM and Hook was used to obtain a cMED-HookΔ map at an average resolution of 4.7 Å. Multiple runs of 3D classification were used to obtain a cMED-CKM class of 6,348 particles, which was subjected to 3D refinement that resulted in a ∼11 Å reconstruction (cMED-CKM). To visualize how CKM binds cMED, a loose mask covering the CKM and Hook was created and used to center the CKM-Hook of 105,401 shiny particles for subsequent multiple runs of local masked 3D classification. This resulted in a class in which CKM and the Hook of cMED were clearly visualized. 3D refinement of this class resulted in a 6.7 Å reconstruction (CKM-Hook). For Head-IDR and Middle-IDR, two different loose masks that cover the Head and Middle, respectively, were created and used to center their corresponding regions of 105,401 shiny particles for performing focused 3D Classification following the similar procedures as described above. 3D refinement of Head-IDR and Middle-IDR classes resulted in two 4.9 Å reconstructions. The image analysis procedures and cryo-EM data collections of CKM and cMED-CKM are shown in Figure S2 and Table S1, respectively.

### Model building and refinement

To build an atomic model of the CKM, we used the x-ray structure of human CDK8/CCNC [Protein Data Bank (PDB ID): 3RGF] and AlphaFold models of human MED12 and MED13 as templates, which were fitted into the cryo-EM maps of the CKM by rigid-body fitting using Chimera^76^. After fitting, some regions of the templates, such as MED12N, MED12C, MED13 IDR, and flexible loops/coils that showed different conformations from the cryo-EM maps or had no density were removed from the model. The regions of the MED12N and MED12C were of sufficient quality for *ab initio* model building. The model building and adjustments were done using Coot^77^. Model refinement of the Kinase-lobe, Kinase/Central-lobes. and H-lobe against their corresponding cryo-EM maps were done by using the real-space refinement in Phenix^78^. These models were fitted into the overall map of CKM and then combined as the final overall structure.

Model building for cMED-CKM was facilitated by using our CKM model and published cryo-EM structure of human cMED [PDB 7EMF] in which we removed MED26 due to its absence in our cMED-CKM specimen. These two structures were firstly fitted into the overall cryo-EM map of cMED-CKM using Chimera, resulting in an overall model of the cMED-CKM complex. The model of CKM with the Hook of MED, including MED10, MED19, and the N-terminus of MED14, from the overall model was then fitted into the cryo-EM map of CKM-Hook, manually adjusted using Coot, and refined by real-space refinement in Phenix. The cMED structure without the Hook was divided into Head, Middle, and Tail portions, and individually fitted into the cryo-EM map of the cMED-HookΔ map by rigid-body fitting using Chimera. The fitted cMED model was manually adjusted and refined using the same procedure as described above. Both refined models of CKM-Hook and cMED without Hook were combined to generate an overall structure of cMED-CKM. The final overall models of CKM and cMED-CKM were validated using MolProbity (table S1)^78^. Phenix was used to calculate real-space map-model difference maps. All molecular graphic figures, including overall and local density maps, were made by Chimera, UCSF Chimera X^79^, or PyMOL^80^.

### Kinase assay

A GST-CTD-6xHis plasmid that contains the CTD domain of human Pol II RPB1 (residues 1593- 1970) with a 6xHis-tag at its C-terminus was expressed in *E. coli* and purified using glutathione Sepharose 4B beads (GE Healthcare). Purified CKM (3 μM) or cMED-CKM (3 μM) was incubated at 30°C for 30 min in a kinase reaction buffer (100 μL), containing 1x PBS (pH 7.4), 10 mM MgCl_2_, 1 mM ATP, and purified GST-CTD-6xHis substrate. Reactions were terminated by adding 20 μL 6xSDS sample buffer. Samples were resolved on 10% SDS-PAGE and analyzed by western blot using anti-Pol II-CTD antibody (8WG16) and monoclonal anti-FLAG M2 antibody (MilliporeSigma, F1804). The antibody (Cell Signaling, 13523) that recognizes phosphorylated Ser5 of CTD was used to detect CTD phosphorylation.

### XL-MS analyses of the CKM and cMED-CKM

The recombinant human CKM purified by anti-Flag M2 affinity beads and glycerol gradient centrifugation was dialyzed into buffer (20 mM Hepes pH 7.9, 150 mM NaCl, and 0.1mM EDTA) and concentrated to ∼ 1 mg/ml. Around 100 ug CKM protein was incubated with disuccinimidyl dibutyric urea crosslinker (DSBU) at a final concentration of 6 mM for 2 hours on ice. The reaction was quenched with Ammonium Bicarbonate and further incubated for 30 min on ice. Crosslinked proteins were subjected to TCA precipitation. For cMED-CKM, proteins eluted from anti-Flag M2 beads were dialyzed into buffer (20 mM Hepes pH7.6, 1 mM MgCl_2_, 200 mM NaCl, 0.25 mM EDTA, and 1 mM beta-mercaptoethanol) and concentrated to ∼1.5 mg/ml. Around 200 ug cMED- CKM protein was crosslinked, quenched, and precipitated following the same procedures applied to CKM.

Crosslinked proteins were reduced with 10 mM DTT for 30 min at 30°C, followed by alkylation with iodoacetamide (50 mM final, Sigma Aldrich) for 30 min at 30°C. The proteins were processed by S-TrapTM (Protifi) with its recommended protocol: with trypsin in 1:10 (w/w) enzyme-to- protein ratio for an hour at 30°C. Eluted peptides were dried under vacuum and resuspended with the peptide fractionation elution buffer: LC-MS grade 70% (v/v) water, 30% (v/v) Acetonitrile (ACN), and 0.1% (v/v) Trifluoroacetic acid (TFA). Peptide fraction was performed on AKTA pure 25 with Superdex 30 Increase 3.2/300 (GE Healthcare) at a flow rate of 30 μl min-1 of the elution buffer with a 100 μl fraction volume. Fractions containing enriched crosslinked peptides, which were empirically determined by the elution profile, were retained and dried under vacuum and resuspended with 0.1% (v/v) TFA containing LC-MS grade water for mass spectrometry analysis. Each fraction was analyzed on a Q-Exactive HF mass spectrometer (Thermo Fisher Scientific) coupled with Dionex Ultimate 3000 UHPLC system (Thermo Fischer Scientific) with in-house C18 column. Half of each sample amount was injected for the analysis and separated on 90-min gradient: mobile phase A (99.9% water with 0.1% formic acid (Sigma Aldrich); mobile phase B (80% ACN with 0.1% formic acid); starting 5% B, increased to 45% B for 90 min, then keep B constant at 90% for 5 min, and sharply decreased to 5% B for 5 min for re-equilibration of the column with the constant flow rate set to 400 nl min-1. The data dependent acquisition method was set as follows: Full MS resolution 120,000; MS1 AGC target 1e6; MS1 Maximum IT 200ms; Scan range 300 to 1800; dd-MS/MS resolution 30,000; MS/MS AGC target 2e5 ; MS2 Maximum IT 300ms; Fragmentation was enforced by higher-energy collisional dissociation with stepped collision energy with 25, 27, 30; Loop count Top 12 ;Isolation window 1.5 ; Fixed first mass 130 ; MS2 Minimum AGC target 800 ; Charge exclusion: unassigned,1,2,3,8,>8 ; Peptide match off ; Exclude isotope on ; Dynamic exclusion 45sec. Raw files were converted to mgf format with TurboRawToMGF 2.0.8^81^: Precursor mass weight range 300 – 10,000 Da and all default removal options were off. Searches for crosslinked peptides were performed by MeroX 2.0.0.5^82^ with the default setting for DSBU with the minor modifications: Mass limit from 300 Da to 10,000 Da, Minimum charge (MS1) set to 4, Apply Prescore and Score cut off to 10, and FDR cut off set to 1%. All search results from each fraction’s MS acquisition was combined and filtered by recalculated FDR at 1%. Redundant crosslinked pairs were sorted by the main score and the top hit was chosen for the final report table and mapping onto the structure in Chimera^76^ with Xlink Analyzer plugin^83^.

### LC-MS-MS analyses of purified Mediator proteins

2 L of 293-F cells expressing Flag-tagged CDK8, MED13 IDRc, or MED26 was used to prepare nuclear extract following the same procedure as described above. Proteins purified from nuclear extract using anti-Flag M2 beads were applied to 15%-40% glycerol gradient centrifugation following the sample protocols as described above. The peak fractions containing large Mediator complexes were merged and dialyzed into buffer (20 mM Hepes pH7.6, 1 mM MgCl_2_, 150 mM NaCl, 1 mM EDTA, 1 mM beta-mercaptoethanol, and 5% glycerol). The protein samples were concentrated and processed by SDS-PAGE. The gel band samples were subjected to In-gel digestion^84^.

An aliquot of the tryptic digest (in 2 % acetonitrile/0.1% formic acid in water) was analyzed by LC/MS/MS on an Orbitrap FusionTM TribridTM mass spectrometer (Thermo ScientificTM) interfaced with a Dionex UltiMate 3000 Binary RSLCnano System. Peptides were separated onto an analytical C18 column (100μm ID x 25 cm, 5 μm, 18Å) at flow rate of 350 nl/min. Gradient conditions were: 3%-22% B for 40 min; 22%-35% B for 10min; 35%-90% B for 10 min; 90% B held for 10 min (solvent A, 0.1 % formic acid in water; solvent B, 0.1% formic acid in acetonitrile). The peptides were analyzed using data-dependent acquisition method, Orbitrap Fusion was operated with measurement of FTMS1 at resolutions 120,000 FWHM, scan range 350-1500 m/z, AGC target 2E5, and maximum injection time of 50 ms; During a maximum 3 second cycle time, the ITMS2 spectra were collected at rapid scan rate mode, with HCD NCE 34, 1.6 m/z isolation window, AGC target 1E4, maximum injection time of 35 ms, and dynamic exclusion was employed for 20 seconds. The raw data files were processed using Thermo ScientificTM Proteome DiscovererTM software version 1.4, spectra were searched against the Uniprot-Homo sapiens database using Sequest. Search results were trimmed to 1% FDR for strict and 5% for relaxed condition using Percolator. For the trypsin, up to two missed cleavages were allowed. MS tolerance was set 10 ppm; MS/MS tolerance 0.8 Da. Carbamidomethylation on cysteine residues was used as fixed modification; oxidation of methione as well as phosphorylation of serine, threonine and tyrosine was set as variable modifications.

---

**Figure S1.**
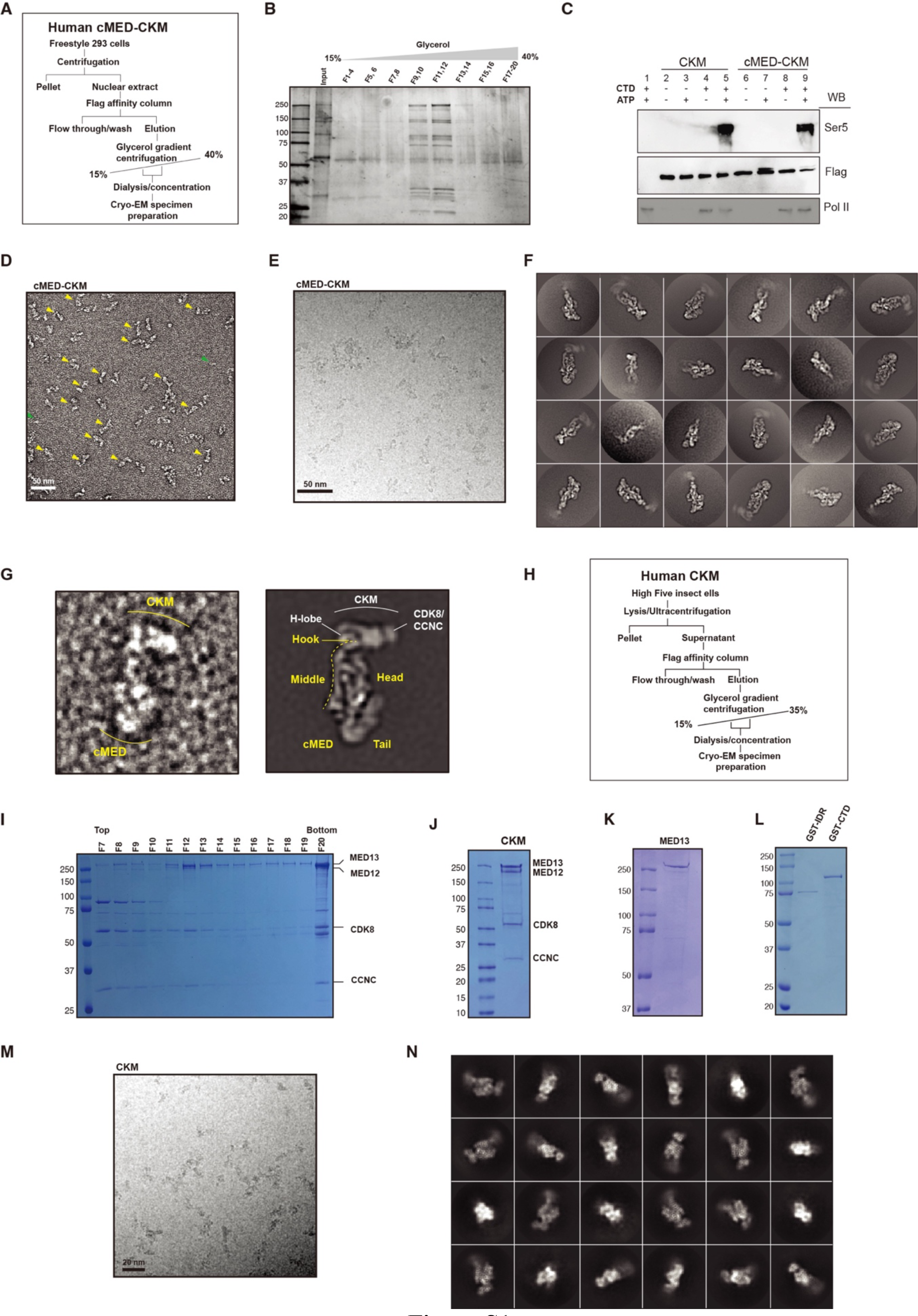
Purification of endogenous human Mediator complex and recombinant CKM. (A) Schematic diagram of two-step purification of the Mediator complex. Flag-tagged CDK8 expressed in 293-F cells was used to purify endogenous cMED-CKM complex. **(B)** Isolation of the cMED-CKM by glycerol gradient centrifugation (15%-40% v/v). Fractions were processed by SDS-PAGE and silver staining. Fractions (F9 to F12) were pooled and subjected to mass spectrometry analysis and structure determination. **(C)** Kinase activity of purified human CKM and cMED-CKM directed toward the Ser5 residue of GST-CTD-6His. Phosphorylated CTD Ser5- specific antibody was used to detect CTD phosphorylation by CDK8. **(D)** A negative-stain micrograph of human cMED-CKM. The particles showing CKM bound to cMED in an approximate perpendicular (∼90 degree) orientation are indicated by yellow arrows. Two CKM- like particles are indicated by green arrows. **(E)** A cryo-EM micrograph showing human cMED- CKM particles preserved in ice. **(F)** Selected cryo-EM 2D averages of the cMED-CKM. **(G)** A raw particle image (left) and 2D class average (right) of the cMED-CKM. **(H)** Schematic diagram of two-step purification of recombinant human CKM expressed in insect cells. **(I)** Isolation of intact CKM by glycerol gradient centrifugation (15%-35% v/v). Gradient fractions were processed by SDS-PAGE and Coomassie blue staining. Fractions (F12 to F15) containing human CKM subunits (CDK8, CCNC, MED12, and MED13) were pooled, dialyzed, and concentrated for subsequent structure determination. **(J)** SDS-PAGE analysis of purified human CKM. **(K)** SDS-PAGE analysis of recombinant human full-length MED13 purified from insect cells. **(L)** SDS-PAGE analysis of GST-CTD and GST-MED13 IDRc expressed and purified from *E.coli*. **(M)** A cryo-EM micrograph showing human CKM particles preserved in ice. **(N)** Representative cryo-EM 2D class averages showing various orientations of the CKM.

**Figure S2.**
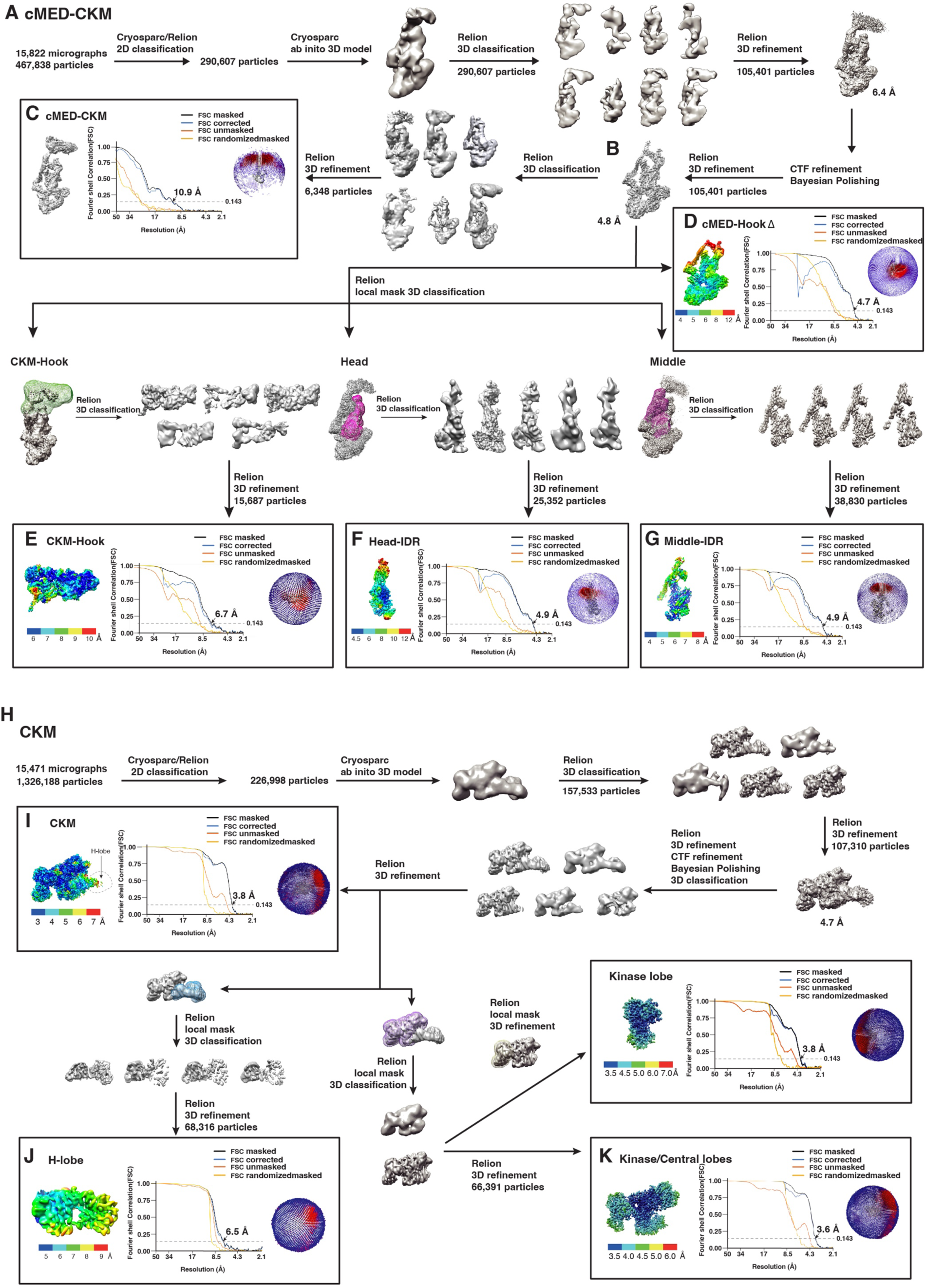
Cryo-EM data processing of the Mediator complex and CKM. (A) Flow-chart for particle selection, 2D/3D classification, and refinement of cMED-CKM cryo-EM images. **(B)** A cMED-CKM cryo-EM map at an average resolution of 4.8 Å after polishing and refinement. **(C)** Left, cryo-EM density map of the cMED-CKM. Middle, FSC curves. Right, angular distribution plot. The resolution of cMED-CKM is 10.9 Å. **(D-G)** Focused refinements of cMED-HookΔ, CKM-Hook, Head-IDR, and Middle-IDR. **(H)** Flow-chart for 2D/3D classification and refinement of CKM cryo-EM images. **(I)** Left, the density map of the entire CKM colored by local resolution. Local resolutions of cryo-EM maps are colored as indicated. Middle, FSC curves. Right, angular distribution plot. The resolution of CKM is 3.8 Å according to the gold standard FSC = 0.143 criterion. **(J,K)** Focused refinements of H-lobe and Kinase/Central-lobes.

**Figure S3.**
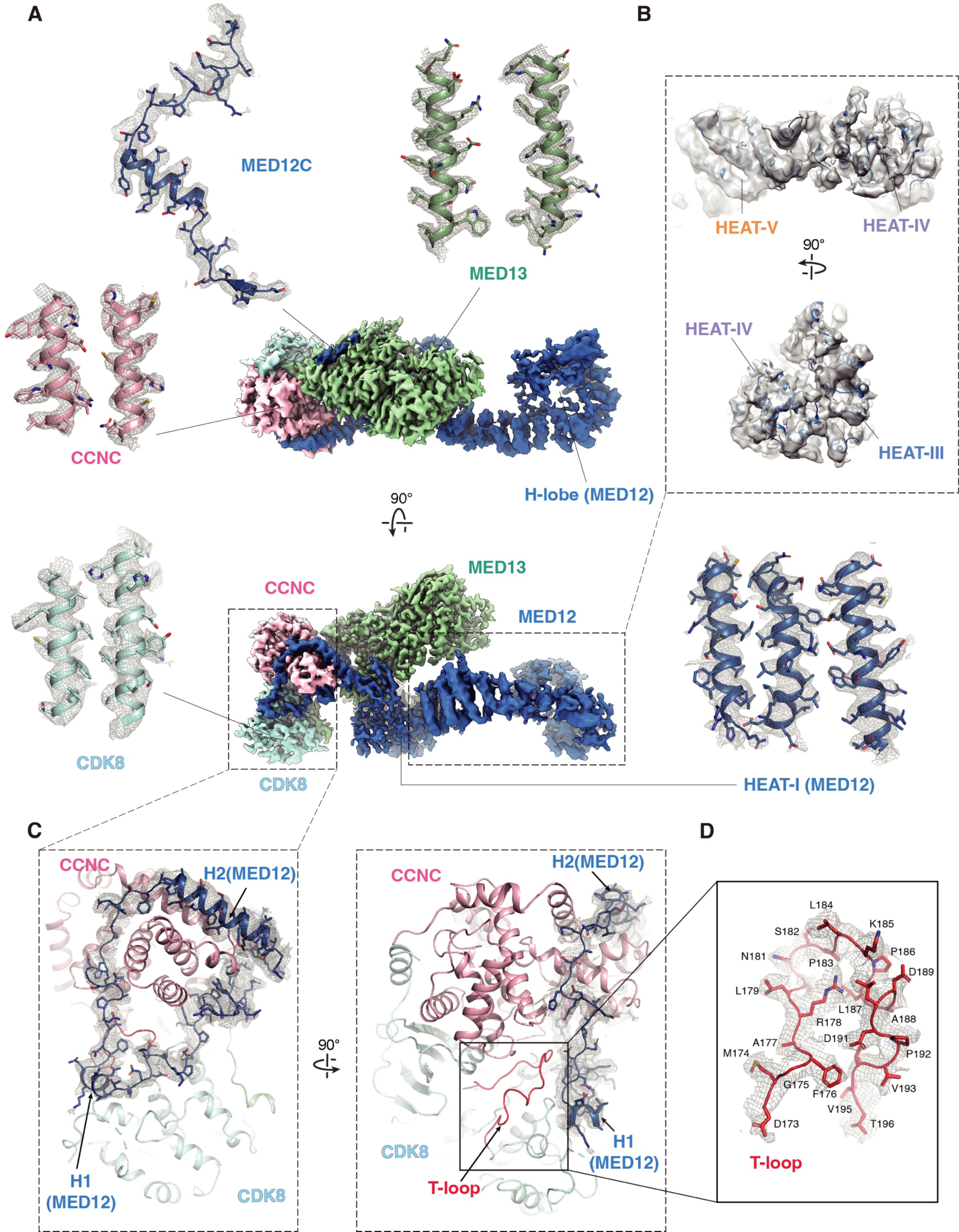
Cryo-EM maps and structural models of human CKM. (A) Two different views of composite cryo-EM map of human CKM. Local structures with their corresponding densities are shown. **(B)** Two close-up views showing density corresponding to the H-lobe that comprises HEAT-II to HEAT-V domains. **(C)** Left, MED12N binds to CDK8/CCNC. MED12N is shown with its corresponding electron density (blue). Right, localization of the T-loop in CDK8 is indicated. **(D)** A close-up view showing the CDK8 T-loop and its corresponding density.

**Figure. S4.**
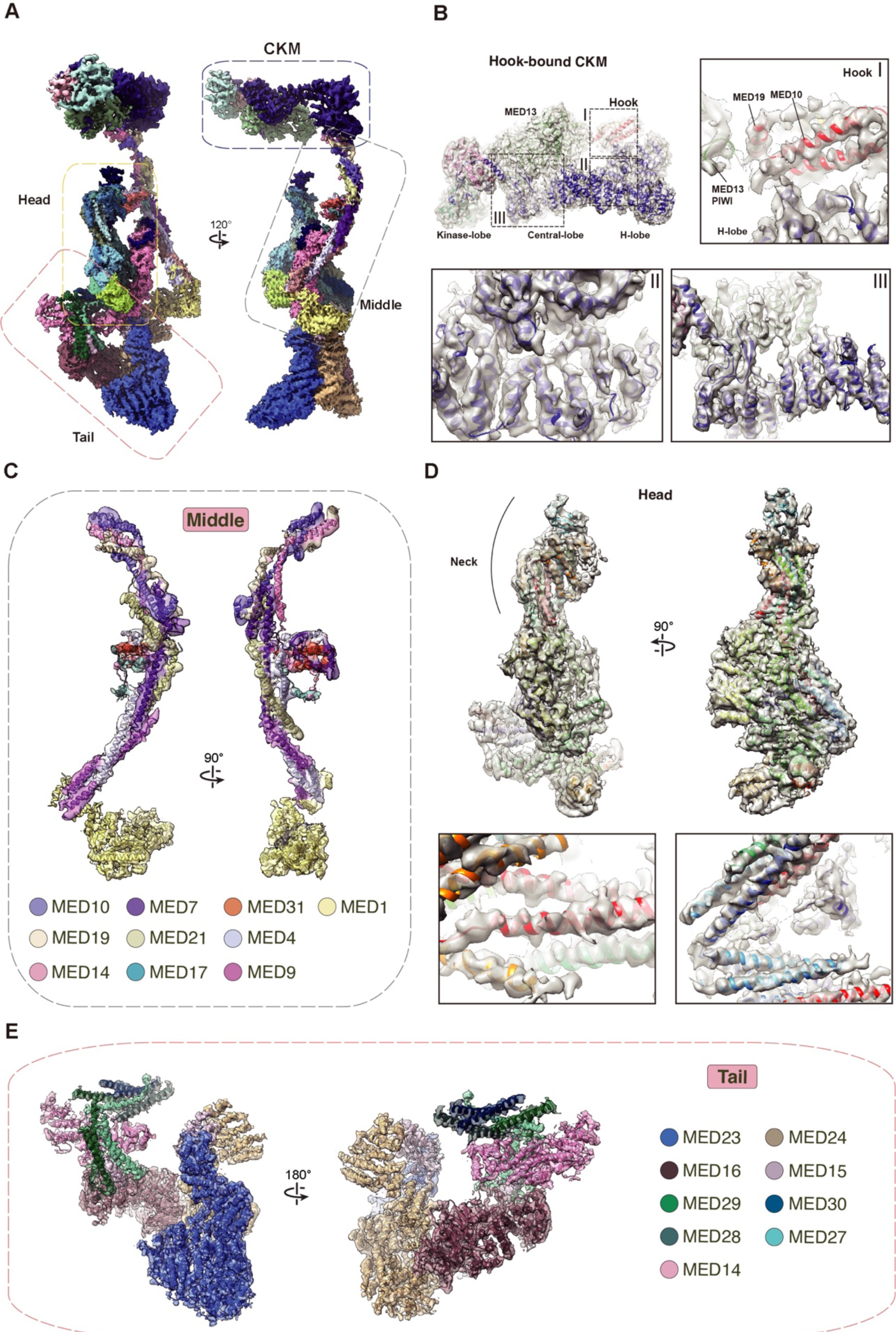
Cryo-EM maps and modular structures of human cMED-CKM. (A) Two different views of the composite cryo-EM map of human cMED-CKM. Head, Middle, and Tail modules as well as the CKM with their corresponding cryo-EM densities are shown in following panels. **(B)** Structure of the CKM bound by the cMED Hook. Local structures with their corresponding cryo-EM densities are shown. **(C)** Structure of the Middle module. The IDR region of MED1 is missing in the density map. **(D)** Structure of the Head module. **(E)** Structure of the Tail module. Each subunit is colored according to color key.

**Figure S5.**
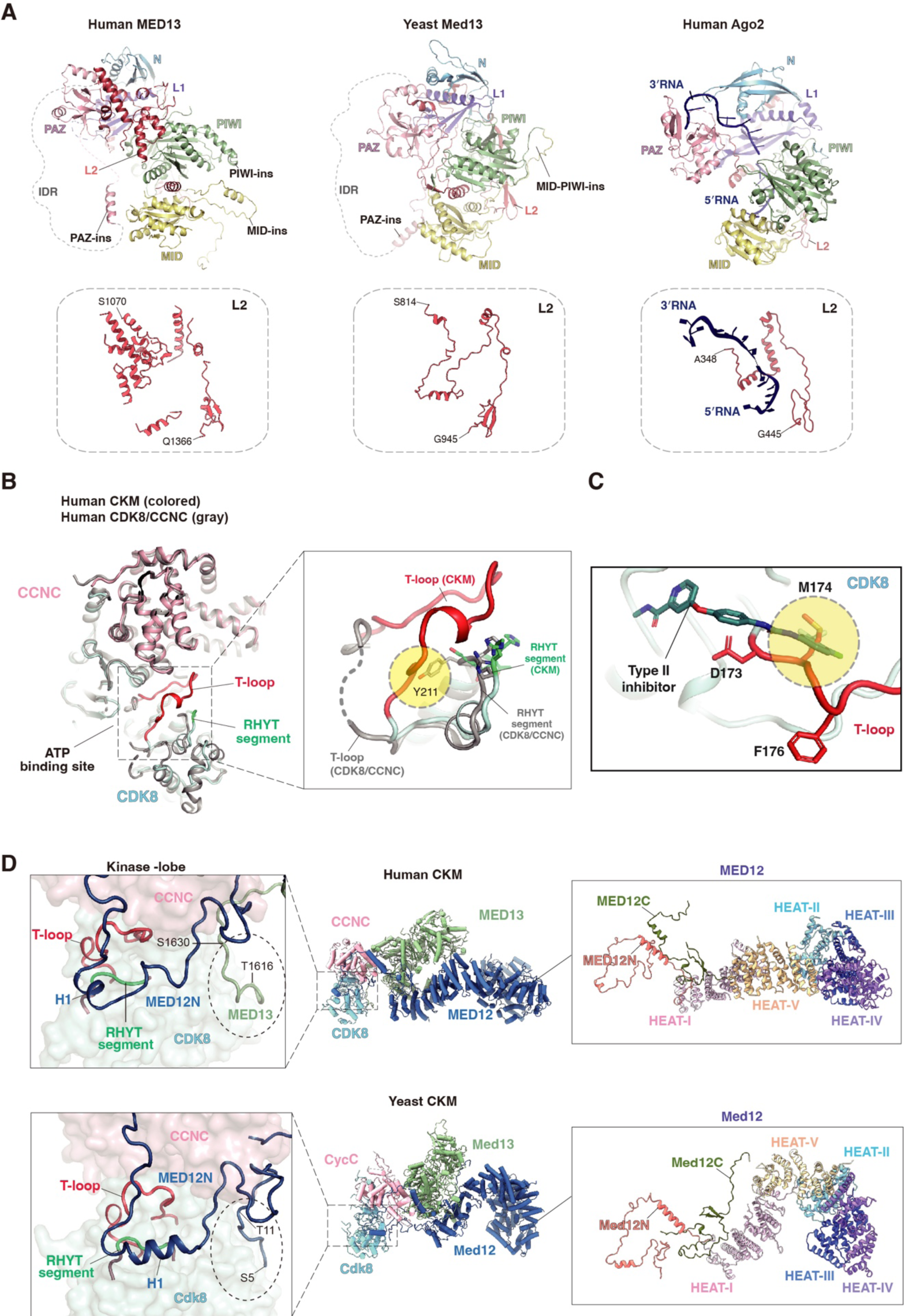
Structural comparison of CKM between human and yeast. (A) Structural comparison of MED13 with human Argonaute-2 (Ago2). Left, human MED13. Middle, yeast Med13 [PDB 7KPX]. Right, human Ago2 [PDB 4W5N]. Their respective L2 domains (red) are shown below. The guide RNA in human Ago2 is colored in blue. The IDRs present in human and yeast MED13 are shown by gray dashed lines. **(B)** Left, a superimposition of the crystal structure of human CDK8/CCNC alone (gray) and the cryo-EM structure of CDK8/CCNC within the CKM. The T-loop in the crystal structure of human CDK8/CCNC without MED12 is missing. Right, a close-up view of the T-loop and RHYT segment. Y211 in the structure of CDK8/CCNC alone that likely clashes with the T-loop is highlighted by a yellow circle. **(C)** The type-II inhibitor: Sorafenib [PDB 3RGF] clashes with the M174 of the DMG motif in CDK8. **(D)**Structural comparison between human and yeast CKM. Left top, both human MED12N and MED13 are involved in CDK8 binding. Left bottom, yeast Cdk8 is bound only by Med12N, but not Med13. In yeast, residues 5-11 of Med12N that contact Cdk8 are replaced by the residues 1616-1630 of MED13 in human (dashed circles). Right, both human and yeast MED12 contain five HEAT domains, but they show distinct conformations.

**Figure S6.**
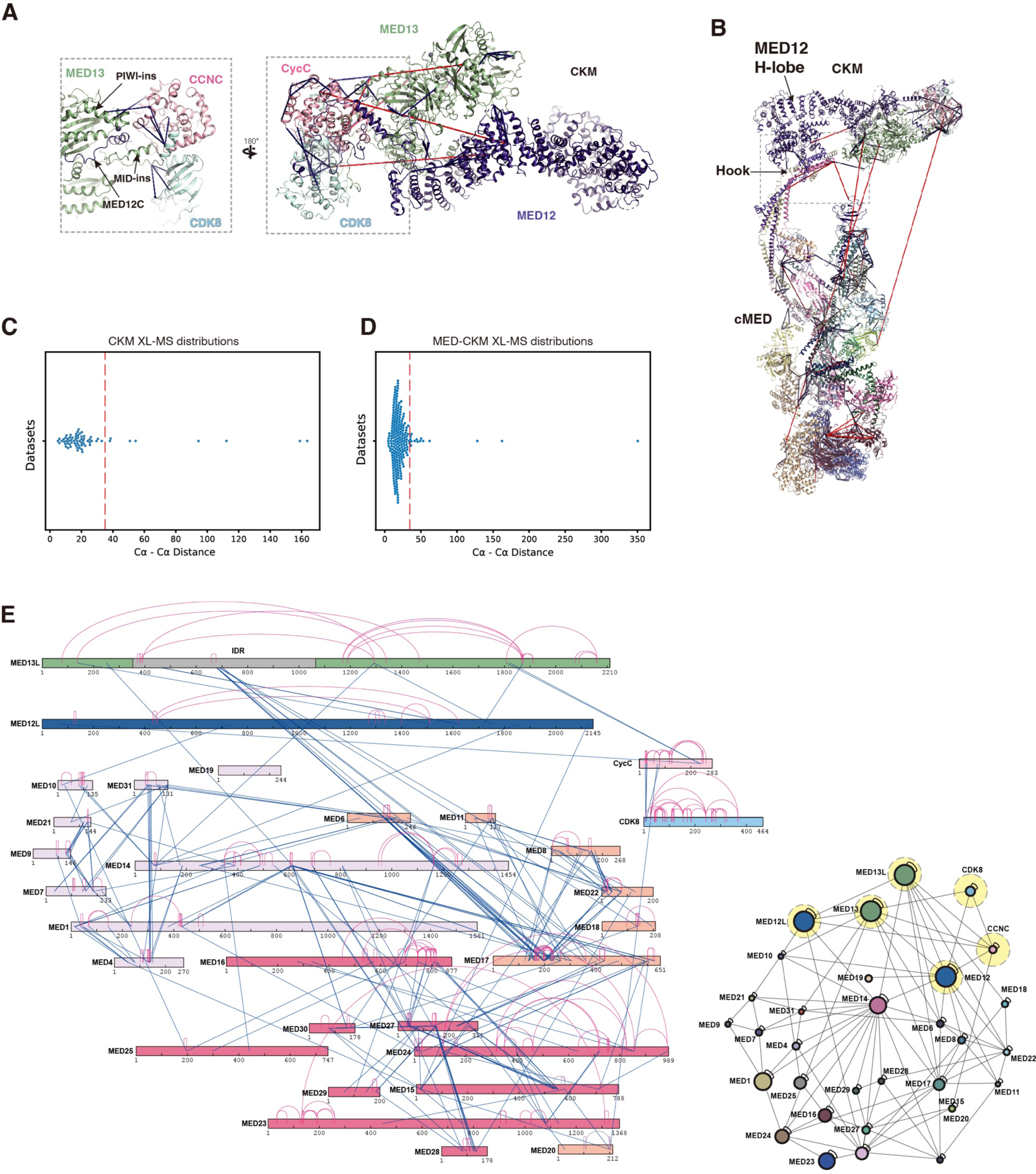
Crosslinking mass spectrometry (XL-MS) analyses of human Mediator complex and its CKM. (A) Mapping all 69 identified URPs onto the CKM. Crosslinks with distances shorter or longer than 35 Å between two Cα-atoms of Lys residues are shown by blue and red lines, respectively. **(B)** Mapping all identified intersubunit and intrasubunit URPs onto the cMED-CKM structure. **(C)** Chart showing the number of cross-links and distances between the Cα atoms of identified URPs of Lysine residues mapped onto the CKM structure with a relatively low violation rate of 13.1 %. **(D)** Chart showing the number of cross-links and distances between the Cα atoms of identified URPs of Lysine residues mapped onto the cMED- CKM structure. In total, 277 URPs were mapped onto the structure with a low violation rate of 8.2 %. **(E)** Crosslinking map of human cMED-CKM, which includes MED12L and MED13L. Intrasubunit and intersubunit URPs identified between lysine residues are shown by red and blue lines, respectively.

**Figure S7.**
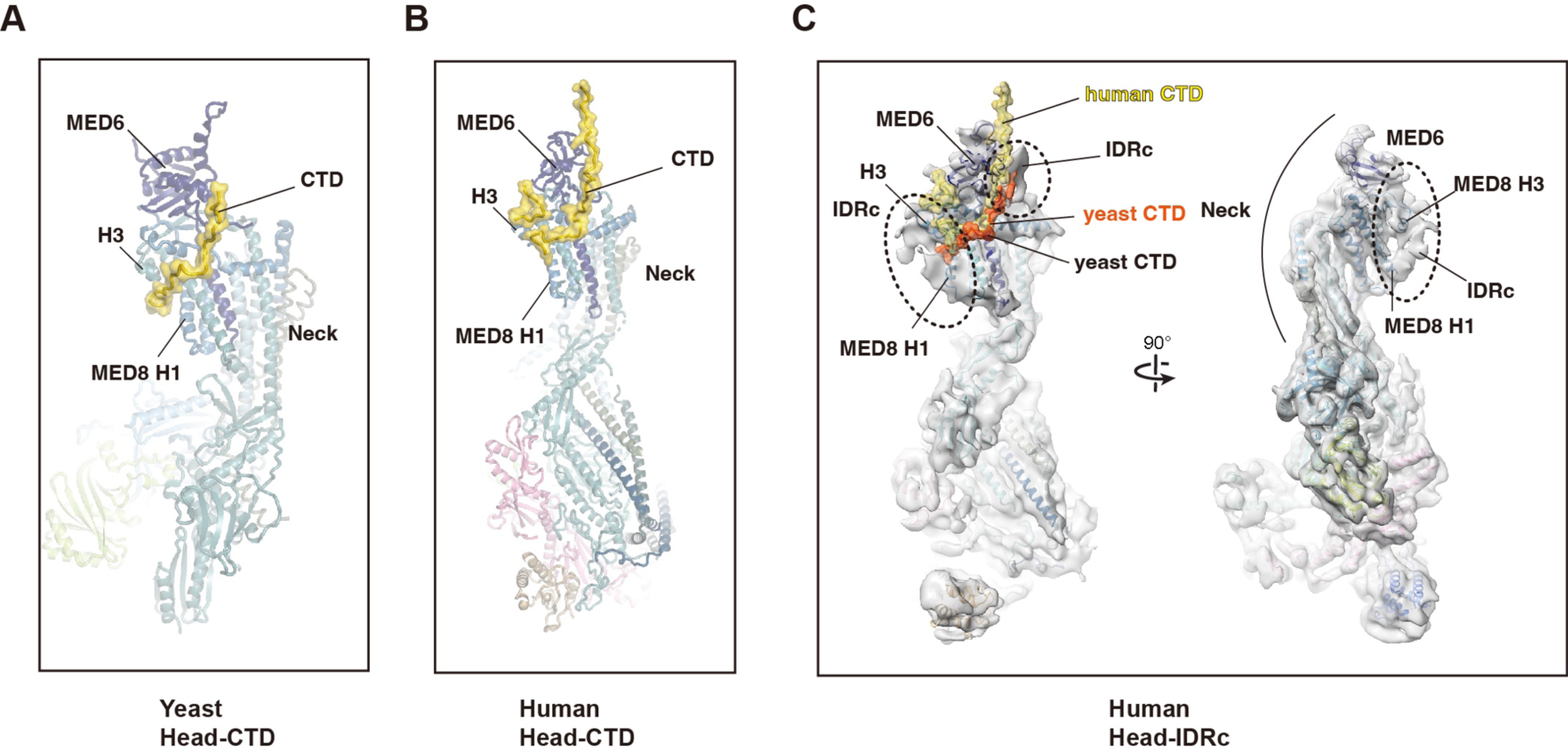
Putative MED13 IDRc densities at Pol II-CTD and MED26 binding sites. (A) Structure of yeast Head-CTD complex [PDB 4GWP]. **(B)** CTD-bound Head module within human cMED-PIC structure [PDB 7ENC]. **(C)** Two different views showing the Head module and its corresponding density map (low-pass filtered to 7Å) within cMED-CKM. Two MED13 IDRc densities next to MED6 and MED8 are indicated by dashed circles. The CTD (orange) of the yeast Head-CTD complex was modeled on the structure showing that its N- and C-terminal regions overlap with MED13 IDRc densities next to MED6 and MED8, respectively. The CTD (yellow) within the human cMED-PIC also overlaps with a MED13 IDRc density next to MED6.

**Figure S8.**
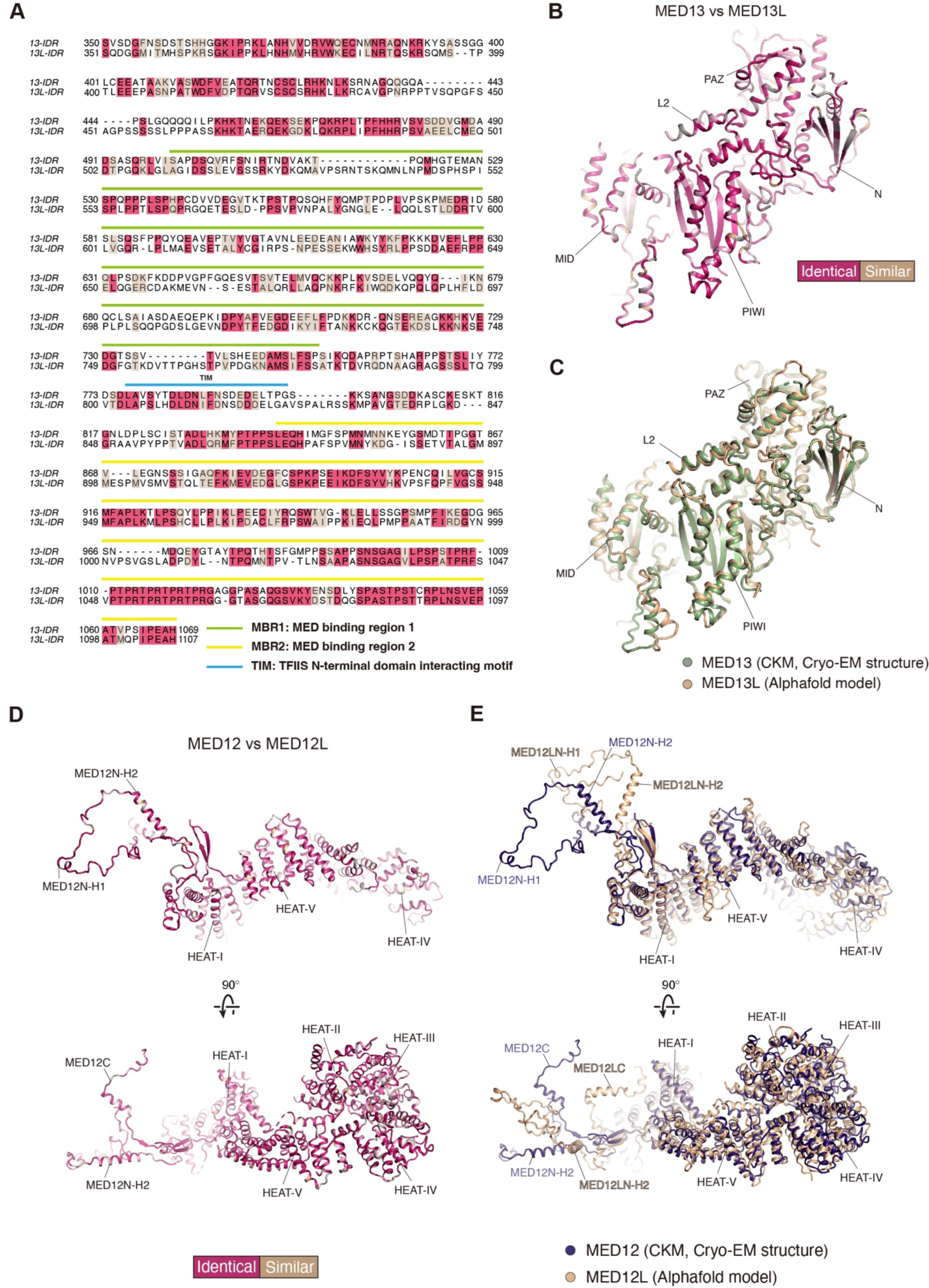
Structural and sequence comparisons of human MED12/12L and MED13/13L. **(A)** Sequence alignment of the IDR regions in human MED13 and MED13L. Identical and similar amino acids are highlighted in red and wheat, respectively. Regions corresponding to MBR1, MBR2, and TIM are as indicated. **(B)** The structure of MED13 is colored according to sequence similarity (67.0% full-length) between MED13 and MED13L. The IDR region is missing in the structure. **(C)** Superimposition of the cryo-EM structure of MED13 (green) and the AlphaFold model of MED13L (brown). The cryo-EM structure of MED13 is similar to the AlphaFold model of MED13L with a Cα RMSD of 1.02 Å (calculated using structured regions). The unstructured regions of MED13L in the AlphaFold model are removed for clarity. **(D)** The MED12 structure is colored according to sequence similarity (71.5% full-length) between MED12 and MED12L. **(E)** Superimposition of the cryo-EM structure of MED12 (blue) and the AlphaFold model of MED12L (brown). Excluding MED12N and MED12C, the overall architecture of the HEAT domains between MED12 and MED12L are similar with a Cα RMSD of 3.71Å.

**Figure S9.**
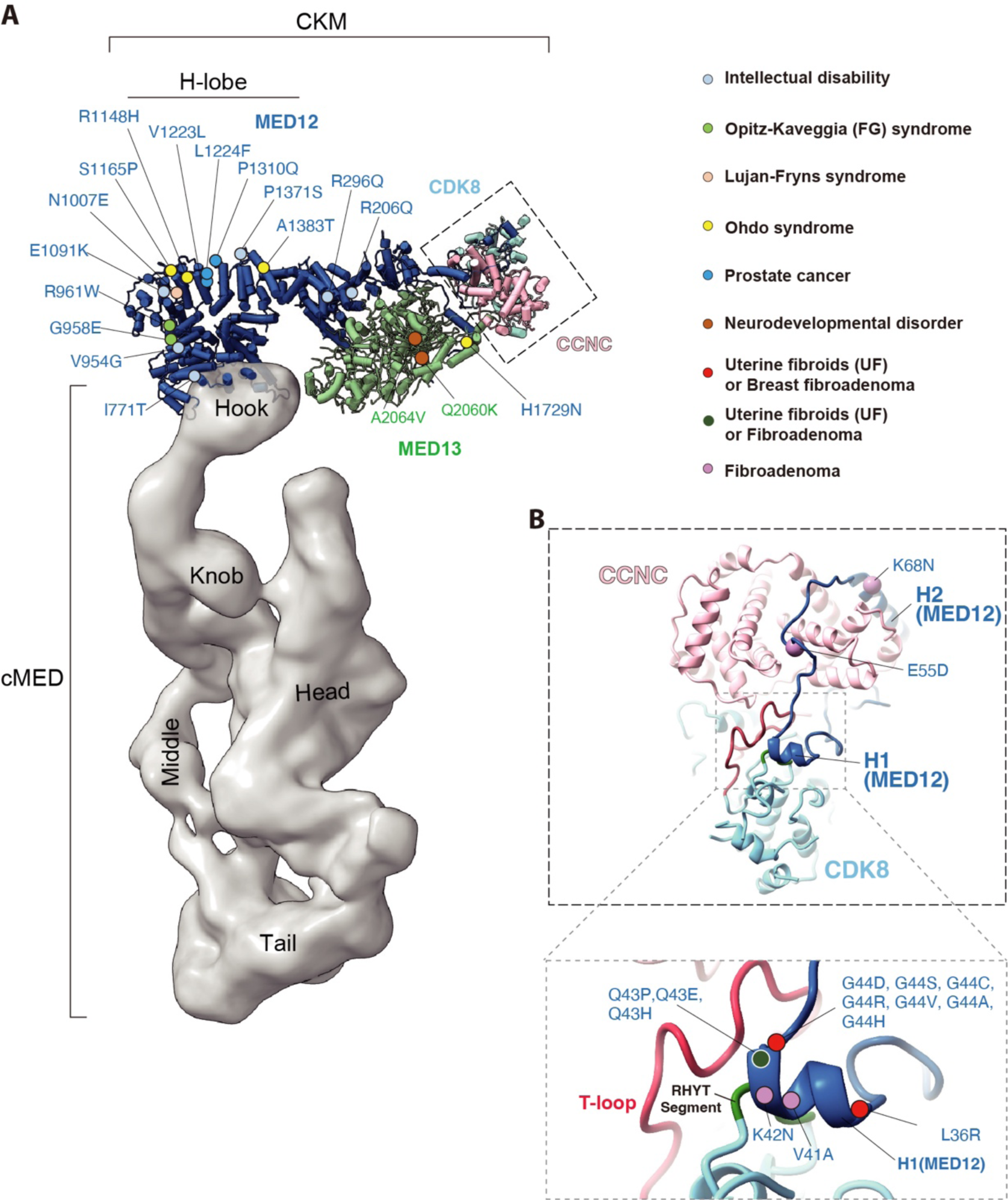
CKM-related disease mutations mapped onto human Mediator structure. (A) The CKM and cMED structure are shown in cylinders and gray surface representation, respectively. Disease mutations are indicated and colored according to the color key shown at right. Numerous MED12 disease mutations are present in the H-lobe. **(B)** Uterine fibroid driver mutations in MED12 are highly clustered in the H1 region (blue) next to the CDK8 T-loop and RHYT segment. Note that the most frequently mutated MED12 residues found in uterine fibroids, Q43 and G44, lie at the interface of the RYHT segment and T-loop in CDK8, consistent with biochemical analyses showing that mutation of these MED12 residues disrupt CDK8 kinase activity.

**Table S1.** Data collection, processing, and refinement statistics.

| Cryo-EM Data Collection | CKM |  |  |  | cMED-CKM |  |  |  |
| --- | --- | --- | --- | --- | --- | --- | --- | --- |
| Microscope | Titan Krios G3 |  |  |  | Titan Krios G3 |  |  |  |
| Voltage (keV) | 300 |  |  |  | 300 |  |  |  |
| Camera | K2 GIF |  |  |  | K2 GIF |  |  |  |
| Defocus range (um) | 0.6-3.0 |  |  |  | 0.6-2.8 |  |  |  |
| Pixel size (Å) | 1.07 |  |  |  | 1.42 |  |  |  |
| Movies | 15,471 |  |  |  | 15,822 |  |  |  |
| Frames per movie | 40 |  |  |  | 48 |  |  |  |
| Exposure time | 8 |  |  |  | 12 |  |  |  |
| Magnification | 130,000 |  |  |  | 105,000 |  |  |  |
| Total dose per movie (e-/Å²) | 64 |  |  |  | 52 |  |  |  |
| Cryo-EM image Processing |  |  |  |  |  |  |  |  |
|  | CKM |  |  |  | cMED-CKM |  |  |  |
| Map | CKM | H-lobe | Kinase lobe | Kinase/ Central-lobes | cMED-CKM | CKM-Hook | Head-IDR | Middle-IDR |
| Particles in the final map | 122,015 | 68,316 | 66,391 | 66,391 | 6,348 | 15,687 | 25,352 | 38,830 |
| Symmetry imposed | C1 | C1 | C1 | C1 | C1 | C1 | C1 | C1 |
| Map resolution (Å) | 3.8 | 6.5 | 3.8 | 3.6 | 10.9 | 6.7 | 4.9 | 4.9 |
| B factor (Å²) | -114 | -714 | -116 | -115 | -437 | -160 | -145 | -122 |
| Model Building & Refinement |  |  |  |  |  |  |  |  |
| Models | CKM | H-lobe | Kinase lobe | Kinase/Central lobes | cMED-CKM | CKM-Hook | Head-IDR | Middle-IDR |
| Initial Model used (PDB) | 3RGF | AlphaFold | 3RGF | 3RGF | 3RGF | 3RGF | 7EMF | 7EMF |
|  |  |  |  |  | 7EMF | 7EMF |  |  |
| Metals | Zn | N/A | N/A | Zn | Zn | Zn | Zn | Zn |
| Other atoms | N/A | N/A | N/A | N/A | N/A | N/A | N/A | N/A |
| R.m.s. deviations |  |  |  |  |  |  |  |  |
| Bond angles (°) | 0.003 | 0.004 | 0.003 | 0.003 | 0.004 | 0.003 | 0.004 | 0.004 |
| Bond lengths (Å) | 0.567 | 0.999 | 0.553 | 0.518 | 0.948 | 0.885 | 0.913 | 0.953 |
| Ramachandran plot |  |  |  |  |  |  |  |  |
| Favored | 97.7 | 94.19 | 97.62 | 97.10 | 94.29 | 94.81 | 94.48 | 93.79 |
| Outliers | 0.00 | 0.00 | 0.00 | 0.05 | 0.11 | 0.06 | 0.08 | 0.08 |
| Validation |  |  |  |  |  |  |  |  |
| Clash score | 8.27 | 9.29 | 5.85 | 5.28 | 9.40 | 8.48 | 9.55 | 8.98 |
| MolProbity score | 1.51 | 1.73 | 1.40 | 1.44 | 1.89 | 1.82 | 1.88 | 1.89 |

## References

1 Kornberg, R. D. Mediator and the mechanism of transcriptional activation. Trends Biochem Sci 30, 235–239, doi:10.1016/j.tibs.2005.03.011 (2005).

2 Malik, S. & Roeder, R. G. The metazoan Mediator co-activator complex as an integrative hub for transcriptional regulation. Nat Rev Genet 11, 761–772, doi:10.1038/nrg2901 (2010).

3 Soutourina, J. Transcription regulation by the Mediator complex. Nat Rev Mol Cell Biol 19, 262–274, doi:10.1038/nrm.2017.115 (2018).

4 Adelman, K. & Lis, J. T. Promoter-proximal pausing of RNA polymerase II: emerging roles in metazoans. Nat Rev Genet 13, 720–731, doi:10.1038/nrg3293 (2012).

5 Core, L. & Adelman, K. Promoter-proximal pausing of RNA polymerase II: a nexus of gene regulation. Genes Dev 33, 960–982, doi:10.1101/gad.325142.119 (2019).

6 Taatjes, D. J., Naar, A. M., Andel, F., 3rd, Nogales, E. & Tjian, R. Structure, function, and activator-induced conformations of the CRSP coactivator. Science 295, 1058–1062, doi:10.1126/science.1065249 (2002).

7 Knuesel, M. T., Meyer, K. D., Bernecky, C. & Taatjes, D. J. The human CDK8 subcomplex is a molecular switch that controls Mediator coactivator function. Genes Dev 23, 439–451, doi:10.1101/gad.1767009 (2009).

8 Tsai, K. L. et al. A conserved Mediator-CDK8 kinase module association regulates Mediator-RNA polymerase II interaction. Nat Struct Mol Biol 20, 611–619, doi:10.1038/nsmb.2549 (2013).

9 Allen, B. L. & Taatjes, D. J. The Mediator complex: a central integrator of transcription. Nat Rev Mol Cell Biol 16, 155–166, doi:10.1038/nrm3951 (2015).

10 Conaway, R. C. & Conaway, J. W. The Mediator complex and transcription elongation. Biochim Biophys Acta 1829, 69–75, doi:10.1016/j.bbagrm.2012.08.017 (2013).

11 Takahashi, H. et al. Human mediator subunit MED26 functions as a docking site for transcription elongation factors. Cell 146, 92–104, doi:10.1016/j.cell.2011.06.005 (2011).

12 Hengartner, C. J. et al. Association of an activator with an RNA polymerase II holoenzyme. Genes Dev 9, 897–910, doi:10.1101/gad.9.8.897 (1995).

13 Firestein, R. et al. CDK8 is a colorectal cancer oncogene that regulates beta-catenin activity. Nature 455, 547–551, doi:10.1038/nature07179 (2008).

14 Park, M. J. et al. Oncogenic exon 2 mutations in Mediator subunit MED12 disrupt allosteric activation of cyclin C-CDK8/19. J Biol Chem 293, 4870–4882, doi:10.1074/jbc.RA118.001725 (2018).

15 Klatt, F. et al. A precisely positioned MED12 activation helix stimulates CDK8 kinase activity. Proc Natl Acad Sci U S A 117, 2894–2905, doi:10.1073/pnas.1917635117 (2020).

16 Danette L. Daniels, M. F., Marie K. Schwinn, Hélène Benink, Matthew D. Galbraith, Ravi Amunugama, Richard Jones, David Allen, Noriko Okazaki, Hisashi Yamakawa, Futaba Miki, Takahiro Nagase, Joaquín M. Espinosa, and Marjeta Urh. Mutual Exclusivity of MED12/MED12L, MED13/13L, and CDK8/19 Paralogs Revealed within the CDK- Mediator Kinase Module. Journal of Proteomics & Bioinformatics (2013).

17 Elmlund, H. et al. The cyclin-dependent kinase 8 module sterically blocks Mediator interactions with RNA polymerase II. Proc Natl Acad Sci U S A 103, 15788–15793, doi:10.1073/pnas.0607483103 (2006).

18 Donner, A. J., Ebmeier, C. C., Taatjes, D. J. & Espinosa, J. M. CDK8 is a positive regulator of transcriptional elongation within the serum response network. Nat Struct Mol Biol 17, 194–201, doi:10.1038/nsmb.1752 (2010).

19 Nemet, J., Jelicic, B., Rubelj, I. & Sopta, M. The two faces of Cdk8, a positive/negative regulator of transcription. Biochimie 97, 22–27, doi:10.1016/j.biochi.2013.10.004 (2014).

20 Galbraith, M. D. et al. HIF1A employs CDK8-mediator to stimulate RNAPII elongation in response to hypoxia. Cell 153, 1327–1339, doi:10.1016/j.cell.2013.04.048 (2013).

21 Steinparzer, I. et al. Transcriptional Responses to IFN-gamma Require Mediator Kinase- Dependent Pause Release and Mechanistically Distinct CDK8 and CDK19 Functions. Mol Cell 76, 485–499 e488, doi:10.1016/j.molcel.2019.07.034 (2019).

22 Poss, Z. C. et al. Identification of Mediator Kinase Substrates in Human Cells using Cortistatin A and Quantitative Phosphoproteomics. Cell Rep 15, 436–450, doi:10.1016/j.celrep.2016.03.030 (2016).

23 Barron, L. et al. Identification and characterization of the mediator kinase-dependent myometrial stem cell phosphoproteome. F S Sci 2, 383–395, doi:10.1016/j.xfss.2021.09.003 (2021).

24 Abdella, R. et al. Structure of the human Mediator-bound transcription preinitiation complex. Science 372, 52–56, doi:10.1126/science.abg3074 (2021).

25 Zhao, H. et al. Structure of mammalian Mediator complex reveals Tail module architecture and interaction with a conserved core. Nat Commun 12, 1355, doi:10.1038/s41467-021-21601-w (2021).

26 Chen, X. et al. Structures of the human Mediator and Mediator-bound preinitiation complex. Science 372, doi:10.1126/science.abg0635 (2021).

27 Rengachari, S., Schilbach, S., Aibara, S., Dienemann, C. & Cramer, P. Structure of the human Mediator-RNA polymerase II pre-initiation complex. Nature 594, 129–133, doi:10.1038/s41586-021-03555-7 (2021).

28 Tsai, K. L. et al. Mediator structure and rearrangements required for holoenzyme formation. Nature 544, 196–201, doi:10.1038/nature21393 (2017).

29 Nozawa, K., Schneider, T. R. & Cramer, P. Core Mediator structure at 3.4 A extends model of transcription initiation complex. Nature 545, 248–251, doi:10.1038/nature22328 (2017).

30 El Khattabi, L., et al. A Pliable Mediator Acts as a Functional Rather Than an Architectural Bridge between Promoters and Enhancers. Cell 178, 1145–1158 e1120, doi:10.1016/j.cell.2019.07.011 (2019).

31 Li, Y. C. et al. Structure and noncanonical Cdk8 activation mechanism within an Argonaute-containing Mediator kinase module. Sci Adv 7, doi:10.1126/sciadv.abd4484 (2021).

32 Schneider, E. V. et al. The structure of CDK8/CycC implicates specificity in the CDK/cyclin family and reveals interaction with a deep pocket binder. J Mol Biol 412, 251–266, doi:10.1016/j.jmb.2011.07.020 (2011).

33 Vulto-van Silfhout, A. T., et al. Mutations in MED12 cause X-linked Ohdo syndrome. Am J Hum Genet 92, 401–406, doi:10.1016/j.ajhg.2013.01.007 (2013).

34 Kim, S., Xu, X., Hecht, A. & Boyer, T. G. Mediator is a transducer of Wnt/beta-catenin signaling. J Biol Chem 281, 14066–14075, doi:10.1074/jbc.M602696200 (2006).

35 Ding, N. et al. Mediator links epigenetic silencing of neuronal gene expression with x- linked mental retardation. Mol Cell 31, 347–359, doi:10.1016/j.molcel.2008.05.023 (2008).

36 Osman, S. et al. The Cdk8 kinase module regulates interaction of the mediator complex with RNA polymerase II. J Biol Chem 296, 100734, doi:10.1016/j.jbc.2021.100734 (2021).

37 Dimitrova, E. et al. Distinct roles for CKM-Mediator in controlling Polycomb-dependent chromosomal interactions and priming genes for induction. Nat Struct Mol Biol 29, 1000–1010, doi:10.1038/s41594-022-00840-5 (2022).

38 Davis, M. A. et al. The SCF-Fbw7 ubiquitin ligase degrades MED13 and MED13L and regulates CDK8 module association with Mediator. Genes Dev 27, 151–156, doi:10.1101/gad.207720.112 (2013).

39 Robinson, P. J., Bushnell, D. A., Trnka, M. J., Burlingame, A. L. & Kornberg, R. D. Structure of the mediator head module bound to the carboxy-terminal domain of RNA polymerase II. Proc Natl Acad Sci U S A 109, 17931–17935, doi:10.1073/pnas.1215241109 (2012).

40 Gorbea Colon, J. J., et al. Structural basis of a transcription pre-initiation complex on a divergent promoter. Mol Cell 83, 574–588 e511, doi:10.1016/j.molcel.2023.01.011 (2023).

41 Sato, S. et al. Role for the MED21-MED7 Hinge in Assembly of the Mediator-RNA Polymerase II Holoenzyme. J Biol Chem 291, 26886–26898, doi:10.1074/jbc.M116.756098 (2016).

42 Freitas, K. A. et al. Enhanced T cell effector activity by targeting the Mediator kinase module. Science 378, eabn5647, doi:10.1126/science.abn5647 (2022).

43 Chen, X. et al. Structures of +1 nucleosome-bound PIC-Mediator complex. Science 378, 62–68, doi:10.1126/science.abn8131 (2022).

44 Naar, A. M., Taatjes, D. J., Zhai, W., Nogales, E. & Tjian, R. Human CRSP interacts with RNA polymerase II CTD and adopts a specific CTD-bound conformation. Genes Dev 16, 1339–1344, doi:10.1101/gad.987602 (2002).

45 Richter, W. F., Nayak, S., Iwasa, J. & Taatjes, D. J. The Mediator complex as a master regulator of transcription by RNA polymerase II. Nat Rev Mol Cell Biol 23, 732–749, doi:10.1038/s41580-022-00498-3 (2022).

46 Liao, S. M. et al. A kinase-cyclin pair in the RNA polymerase II holoenzyme. Nature 374, 193–196, doi:10.1038/374193a0 (1995).

47 Sato, S. et al. A set of consensus mammalian mediator subunits identified by multidimensional protein identification technology. Mol Cell 14, 685–691, doi:10.1016/j.molcel.2004.05.006 (2004).

48 Cevher, M. A. et al. Reconstitution of active human core Mediator complex reveals a critical role of the MED14 subunit. Nat Struct Mol Biol 21, 1028–1034, doi:10.1038/nsmb.2914 (2014).

49 Burroughs, A. M., Iyer, L. M. & Aravind, L. Two novel PIWI families: roles in inter- genomic conflicts in bacteria and Mediator-dependent modulation of transcription in eukaryotes. Biol Direct 8, 13, doi:10.1186/1745-6150-8-13 (2013).

50 Cermakova, K. et al. A ubiquitous disordered protein interaction module orchestrates transcription elongation. Science 374, 1113–1121, doi:10.1126/science.abe2913 (2021).

51 Fan, X., Chou, D. M. & Struhl, K. Activator-specific recruitment of Mediator in vivo. Nat Struct Mol Biol 13, 117–120, doi:10.1038/nsmb1049 (2006).

52 Jeronimo, C. et al. Tail and Kinase Modules Differently Regulate Core Mediator Recruitment and Function In Vivo. Mol Cell 64, 455–466, doi:10.1016/j.molcel.2016.09.002 (2016).

53 Petrenko, N. & Struhl, K. Comparison of transcriptional initiation by RNA polymerase II across eukaryotic species. Elife 10, doi:10.7554/eLife.67964 (2021).

54 Petrenko, N., Jin, Y., Wong, K. H. & Struhl, K. Mediator Undergoes a Compositional Change during Transcriptional Activation. Mol Cell 64, 443–454, doi:10.1016/j.molcel.2016.09.015 (2016).

55 Kagey, M. H. et al. Mediator and cohesin connect gene expression and chromatin architecture. Nature 467, 430–435, doi:10.1038/nature09380 (2010).

56 Zhu, X. et al. Genome-wide occupancy profile of mediator and the Srb8-11 module reveals interactions with coding regions. Mol Cell 22, 169–178, doi:10.1016/j.molcel.2006.03.032 (2006).

57 Andrau, J. C. et al. Genome-wide location of the coactivator mediator: Binding without activation and transient Cdk8 interaction on DNA. Mol Cell 22, 179–192, doi:10.1016/j.molcel.2006.03.023 (2006).

58 Liu, Q. et al. The characterization of Mediator 12 and 13 as conditional positive gene regulators in Arabidopsis. Nat Commun 11, 2798, doi:10.1038/s41467-020-16651-5 (2020).

59 Clark, A. D., Oldenbroek, M. & Boyer, T. G. Mediator kinase module and human tumorigenesis. Crit Rev Biochem Mol Biol 50, 393–426, doi:10.3109/10409238.2015.1064854 (2015).

60 Makinen, N. et al. MED12, the mediator complex subunit 12 gene, is mutated at high frequency in uterine leiomyomas. Science 334, 252–255, doi:10.1126/science.1208930 (2011).

61 Turunen, M. et al. Uterine leiomyoma-linked MED12 mutations disrupt mediator-associated CDK activity. Cell Rep 7, 654–660, doi:10.1016/j.celrep.2014.03.047 (2014).

62 Park, M. J. et al. Mediator Kinase Disruption in MED12-Mutant Uterine Fibroids From Hispanic Women of South Texas. J Clin Endocrinol Metab 103, 4283–4292, doi:10.1210/jc.2018-00863 (2018).

63 Lai, F. et al. Activating RNAs associate with Mediator to enhance chromatin architecture and transcription. Nature 494, 497–501, doi:10.1038/nature11884 (2013).

64 Hoeppner, S., Baumli, S. & Cramer, P. Structure of the mediator subunit cyclin C and its implications for CDK8 function. J Mol Biol 350, 833–842, doi:10.1016/j.jmb.2005.05.041 (2005).

65 Schneider, E. V., Bottcher, J., Huber, R., Maskos, K. & Neumann, L. Structure-kinetic relationship study of CDK8/CycC specific compounds. Proc Natl Acad Sci U S A 110, 8081–8086, doi:10.1073/pnas.1305378110 (2013).

66 Erdos, G., Pajkos, M. & Dosztanyi, Z. IUPred3: prediction of protein disorder enhanced with unambiguous experimental annotation and visualization of evolutionary conservation. Nucleic Acids Res 49, W297–W303, doi:10.1093/nar/gkab408 (2021).

67 Chen, X. et al. Structural insights into preinitiation complex assembly on core promoters. Science 372, doi:10.1126/science.aba8490 (2021).

## References

68 Tomomori-Sato, C., Sato, S., Conaway, R. C. & Conaway, J. W. Immunoaffinity purification of protein complexes from Mammalian cells. Methods Mol Biol 977, 273–287, doi:10.1007/978-1-62703-284-1_22 (2013).

69 Park, M. J. et al. Oncogenic exon 2 mutations in Mediator subunit MED12 disrupt allosteric activation of cyclin C-CDK8/19. J Biol Chem 293, 4870–4882, doi:10.1074/jbc.RA118.001725 (2018).

70 Zheng, S. Q. et al. MotionCor2: anisotropic correction of beam-induced motion for improved cryo-electron microscopy. Nat Methods 14, 331–332, doi:10.1038/nmeth.4193 (2017).

71 Zhang, K. Gctf: Real-time CTF determination and correction. J Struct Biol 193, 1–12, doi:10.1016/j.jsb.2015.11.003 (2016).

72 Wagner, T. & Raunser, S. The evolution of SPHIRE-crYOLO particle picking and its application in automated cryo-EM processing workflows. Commun Biol 3, 61, doi:10.1038/s42003-020-0790-y (2020).

73 Punjani, A., Rubinstein, J. L., Fleet, D. J. & Brubaker, M. A. cryoSPARC: algorithms for rapid unsupervised cryo-EM structure determination. Nat Methods 14, 290–296, doi:10.1038/nmeth.4169 (2017).

74 Scheres, S. H. RELION: implementation of a Bayesian approach to cryo-EM structure determination. J Struct Biol 180, 519–530, doi:10.1016/j.jsb.2012.09.006 (2012).

75 Henderson, R. et al. Outcome of the first electron microscopy validation task force meeting. Structure 20, 205–214, doi:10.1016/j.str.2011.12.014 (2012).

76 Pettersen, E. F. et al. UCSF Chimera--a visualization system for exploratory research and analysis. J Comput Chem 25, 1605–1612, doi:10.1002/jcc.20084 (2004).

77 Emsley, P., Lohkamp, B., Scott, W. G. & Cowtan, K. Features and development of Coot. Acta Crystallogr D Biol Crystallogr 66, 486–501, doi:10.1107/S0907444910007493 (2010).

78 Adams, P. D. et al. PHENIX: a comprehensive Python-based system for macromolecular structure solution. Acta Crystallogr D Biol Crystallogr 66, 213–221, doi:10.1107/S0907444909052925 (2010).

79 Pettersen, E. F. et al. UCSF ChimeraX: Structure visualization for researchers, educators, and developers. Protein Sci 30, 70–82, doi:10.1002/pro.3943 (2021).

80. DeLano, W. L. The PyMOL Molecular Graphics System. (2002).

81 Sheng, Q. et al. Preprocessing significantly improves the peptide/protein identification sensitivity of high-resolution isobarically labeled tandem mass spectrometry data. Mol Cell Proteomics 14, 405–417, doi:10.1074/mcp.O114.041376 (2015).

82 Iacobucci, C. et al. A cross-linking/mass spectrometry workflow based on MS-cleavable cross-linkers and the MeroX software for studying protein structures and protein-protein interactions. Nat Protoc 13, 2864–2889, doi:10.1038/s41596-018-0068-8 (2018).

83 Kosinski, J. et al. Xlink Analyzer: software for analysis and visualization of cross-linking data in the context of three-dimensional structures. J Struct Biol 189, 177–183, doi:10.1016/j.jsb.2015.01.014 (2015).

84 Shevchenko, A. et al. Archived polyacrylamide gels as a resource for proteome characterization by mass spectrometry. Electrophoresis 22, 1194–1203, doi:10.1002/1522- 2683()22:6<1194::AID-ELPS1194>3.0.CO;2-A (2001).

